# Persistent stromal reprogramming defines incomplete mucosal healing and predicts therapeutic response in ulcerative colitis

**DOI:** 10.64898/2026.06.09.730670

**Authors:** Mor Hindi Malowany, Yaniv Stein, Rachel Frieman-Sharabi, Oshrat Levi-Galibov, Neta Yanir, Merav Kedmi, Maor Pauker, Manar Matar, Yifat Snir, Noa Tal, Hagar Banai Eran, Yael Weintraub, Sara Morgenstern, Ofra Golani, Inna Goliand, Yosef Addadi, Hadas Keren-Shaul, Iris Dotan, Raanan Shamir, Shalev Itzkovitz, Henit Yanai, Dror S. Shouval, Ruth Scherz-Shouval

## Abstract

Mucosal healing (MH) is the primary therapeutic endpoint in ulcerative colitis (UC), yet frequent relapses suggest it does not reflect complete tissue recovery. To define the basis of this vulnerability, we generated a multimodal atlas of UC integrating single-cell and bulk transcriptomics, Visium HD spatial profiling, and multiplexed imaging across 89 patients. We show that MH represents a distinct biological state marked by persistent stromal remodeling along three axes: emergence of inflammatory fibroblasts, sustained loss of OGN⁺ niche-supporting fibroblasts, and expansion of pericytes with matrix-remodeling features and reduced vascular association. Spatial analyses revealed persistent reorganization of mucosal tissue domains despite apparent clinical remission. Across independent cohorts, baseline inflammatory fibroblast and pericyte signatures robustly predicted non-response to anti-TNF therapy. These findings suggest that patients in MH remain in a biologically altered state linked to relapse risk and identify stromal reprogramming as a determinant of disease persistence and therapeutic response in UC.

## Introduction

Fibroblasts are stromal cells increasingly recognized as dynamic regulators of tissue homeostasis, inflammation, and tumor progression ^1-6^. In the colon, where the mucosa is exposed to cycles of injury and repair, fibroblasts play a central role in shaping the local microenvironment ^7-11^. Their activation contributes to chronic inflammation and cancer. However, the dynamics and reversibility of these stromal changes remain poorly understood. In particular, it is unclear whether activated stromal cells return to a quiescent state upon clinical remission or whether long-lasting alterations in their activation state persist and contribute to recurrent flares and relapse. Resolving this question is essential for understanding and ultimately targeting the stromal contribution to intestinal diseases. It may also inform broader principles of stromal plasticity in chronic inflammatory disorders and inflammation-associated cancers.

Chronic intestinal inflammation most commonly manifests as inflammatory bowel disease (IBD), encompassing Crohn’s disease (CD) and ulcerative colitis (UC). These chronic relapsing/remitting disorders affect millions of people worldwide and remain a major cause of morbidity ^12,13^. Patients with UC typically present with abdominal pain and bloody diarrhea, resulting from intestinal inflammation confined to the colon ^12-14^. Various therapies are used to induce and maintain remission in patients with UC, with a goal of reaching mucosal healing (MH)^15^, defined as lack of endoscopic inflammation (i.e. endoscopic remission). Despite this, many patients experience repeated flares, and some display a severe, treatment-refractory course that may result in colectomy ^16,17^. Patients with UC are also at increased risk of developing an aggressive form of colorectal cancer termed colitis-associated cancer ^18,19^, necessitating long-term colonoscopic surveillance. The mechanisms underlying the marked heterogeneity in disease course and long-term outcomes in UC remain incompletely understood ^20,21^. Emerging evidence suggests that MH is not fully equivalent to restoration of tissue homeostasis, as residual immune activation and persistent transcriptional alterations have been detected even in patients in endoscopic remission ^22^. However, while immune mechanisms driving inflammation and remission have been extensively studied ^23,24^, the contribution of the stromal microenvironment to the heterogeneous course of MH remains largely unexplored. This raises the possibility that MH represents a clinically defined but biologically altered state, in which residual stromal reprogramming, and potentially a form of stromal “inflammatory memory”, predispose tissues to states of relapse and disease progression.

The intestinal stroma comprises diverse cell populations, including fibroblasts, myofibroblasts, pericytes, smooth muscle cells, and mesenchymal stromal cells which actively promote inflammation and cancer through regulation of the immune microenvironment ^25-27^ and extracellular matrix (ECM) remodeling ^7,9,28-30^. In CD, ECM remodeling contributes to stricturing, fibrosis, and fistula formation ^8,31-33^. In UC, the dynamics of stromal activation and ECM remodeling, and their potential reversibility during MH, remain poorly characterized. In previous work using a mouse model of colitis-associated cancer, we showed that fibroblasts undergo early transcriptional activation during inflammation, and that this activation shapes the trajectory of colitis and cancer ^10^. In human disease, single-cell RNA sequencing (scRNA-seq) studies of intestinal biopsies from patients with UC have identified inflammation-associated fibroblast subsets expressing IL33, TNFSF14, fibroblastic reticular cell markers, and genes linked to epithelial dysfunction and inflammation ^7^. A different study described inflammation-associated fibroblasts (IAFs) that expressed canonical colon cancer-associated fibroblast (CAF) markers such as FAP and WNT2 ^9^, suggesting similarities between stromal programs activated in inflammation and cancer. However, whether these programs are driven by the same cells undergoing dynamic rewiring or by distinct stromal subsets with specialized functions remains unknown. Critically, whether such pathogenic states persist after resolution of inflammation during MH has not been systematically investigated.

Here, we set out to elucidate the cellular and molecular programs that govern stromal activation, persistence, and diversification across inflammation and healing in UC. To this end, we assembled a deeply phenotyped, multi-modal dataset from colonic samples of 89 individuals, spanning healthy controls, active UC, and patients with UC that exhibit MH. By integrating single-cell and bulk transcriptomic analyses with spatially resolved profiling and ECM imaging, we uncovered marked stromal reprogramming during active inflammation and identified stromal states that fail to normalize upon healing. Active disease was characterized by expansion of inflammatory fibroblast subsets and activated pericytes. In parallel, fibroblasts supporting the epithelial stem-cell niche were disrupted during inflammation and only partially recovered during healing, coinciding with incomplete restoration of crypt architecture. Together, these findings demonstrate that MH represents a biologically distinct state marked by persistent, spatially organized stromal reprogramming, positioning the stroma as a central determinant of incomplete healing and relapse susceptibility in UC.

## Results

### scRNA-seq of UC biopsies reveals shifts in epithelial architecture and stromal signaling that are not restored during MH

To comprehensively map the dynamic shift in stromal and epithelial composition during the course of UC, we assembled a cohort of 89 patients representing distinct disease states: healthy controls, active UC (displaying a Mayo Endoscopic Score (MES) ≥1), and UC in MH (MES= 0; Supplementary Table 1). The cohort included both pediatric and adult patients (59 pediatric, 30 adult), reflecting largely conserved mucosal gene expression programs in IBD^34,35^, and enabling analysis across a broad spectrum of UC disease states. Rectal biopsies were collected from all participants together with detailed clinical and demographic data (Supplementary Table 1). From these biopsies, we generated four complementary datasets (Figure 1A). We performed scRNA-seq on a subset of 15 patients to resolve cellular heterogeneity, generated bulk RNA-seq profiles from 43 patients to validate transcriptional signatures, applied Visium HD spatial transcriptomics to samples from 12 patients to obtain high-resolution maps of gene expression within intact tissue architecture, and validated transcriptomic findings using second harmonic generation (SHG) imaging and immunostaining of biopsies from 68 patients (Supplementary Table 1; some patients contributed samples to multiple modalities).

**Figure 1.**
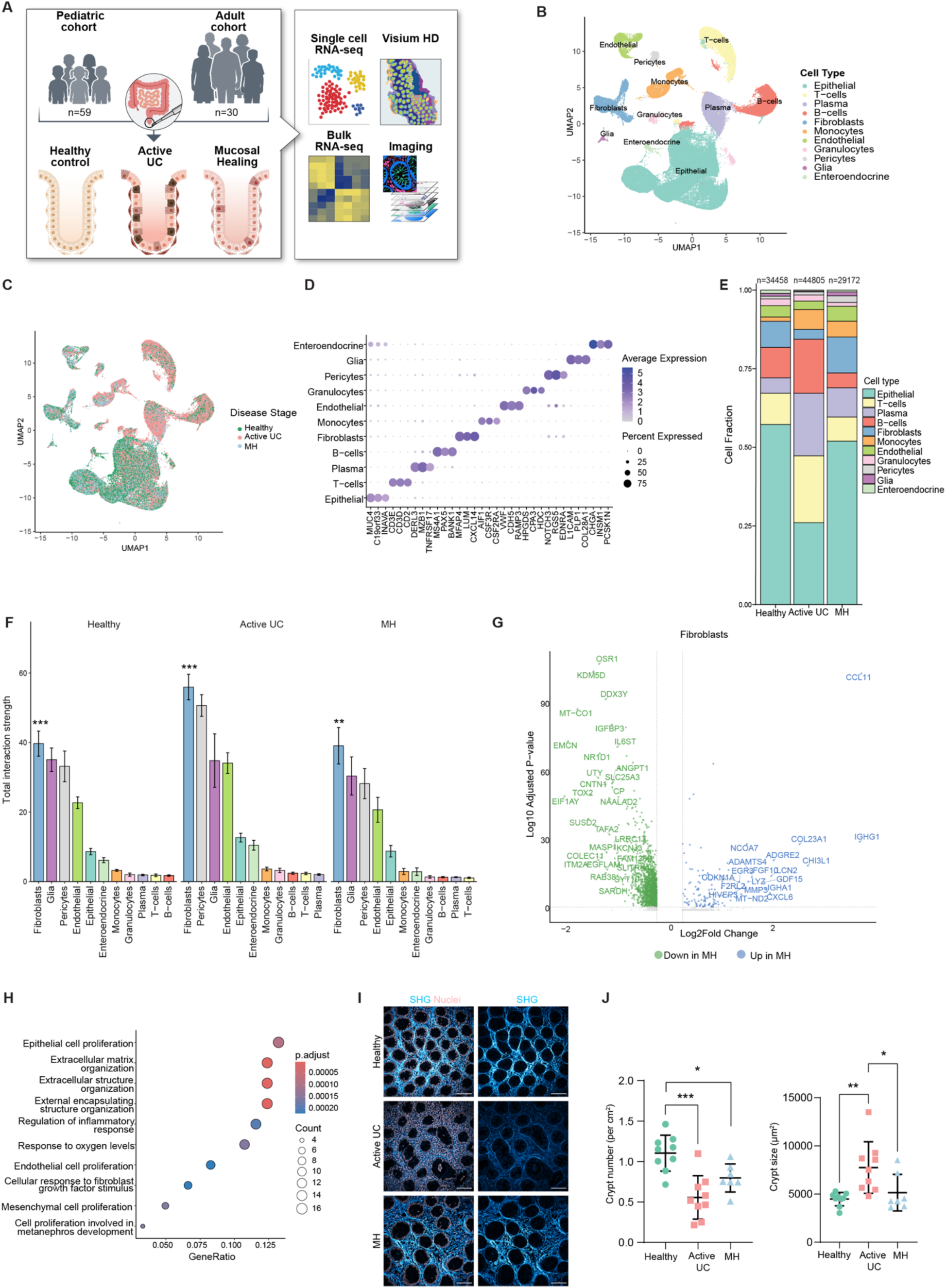
scRNA-seq of UC patients reveals compositional and structural alterations that fail to resolve during MH. **(A)** Schematic overview of the study design. Colonic biopsies were collected from a cohort of pediatric (n = 59) and adult (n = 30) individuals, including healthy controls, patients with active UC, and patients in MH. Tissue samples were processed for scRNA-seq (n=15), bulk RNA-seq (n=43), Visium HD spatial transcriptomics (n=12) and imaging (n=68). **(B-C)** UMAP visualization of 112,435 single cells integrated from 15 patients and analyzed by Seurat, colored by major cell lineage **(B)** and disease state **(C)**. **(D)** Dot plot displaying the top three uniquely expressed genes (UEGs) for each cell lineage, defined using the U-method algorithm. **(E)** Stacked bar plot representation of the relative proportion of cell lineages across disease states (healthy, active UC, and MH). **(F)** Bar plot representation of the signaling activity of each cell lineage, based on cumulative ligand-receptor interaction scores, calculated using the CellChat algorithm applied to the scRNA-seq data described in **(B).** Sender signals are shown. Error bars represent mean ± SEM. Statistical significance between fibroblasts and all other cell lineages within each disease stage was determined using two-sided Wilcoxon rank-sum tests (**p < 0.01, ***p < 0.001). **(G)** Volcano plot displaying differentially expressed genes (DEGs) in the fibroblast cluster of MH patients compared to healthy controls. Significantly upregulated (blue) and downregulated (green) genes in MH are highlighted. Highly significant genes are labeled (Fold Change ≥ 2.5 or ≤0.5, adjusted P≤ 0.005).. **(H)** Top 10 significantly enriched pathways for genes upregulated in the MH fibroblasts, analyzed using clusterProfiler (adjusted Pvalue<0.05). **(I)** Representative second harmonic generation (SHG) images of fibrillar collagen (Cyan), with and without nuclei staining (DRAQ5; light orange), of rectal biopsies from healthy controls, patients with active UC and patients with UC in MH. Scale bar - 100µm **(J)** Quantification of SHG images, including crypt number (left) and crypt size (right), from a cohort of patients including n=9 healthy controls, n=9 active UC and n=7 MH. Results are shown as mean ± SEM analyzed by One-way ANOVA and Tuckey correction for multiple comparisons.

ScRNA-seq was performed on biopsies from 5 healthy controls, 6 patients with active UC, and 4 patients with MH, yielding a total of 112,435 high-quality cells. Seurat clustering followed by UMAP visualization revealed 11 distinct clusters, found across disease states (Figure 1B-C). To define cluster-specific gene signatures and assign cell-type identities we applied a probability-based algorithm which we recently developed termed the U-method ^36^ (https://github.com/YanuvS-Dev/Umethod). The U-method identifies uniquely expressed genes (UEGs) by comparing a gene’s expression probability within a cluster with its highest expression probability across all other clusters. Applying the U-method enabled assignment of canonical cell-type identities (Figure 1D, Supplementary Figure 1A and Supplementary Table 2). Using these annotations, we assessed dynamic changes in cell compositions across disease states (Figure 1E and Supplementary Figure 1B). Analysis of relative cell proportions revealed significant compositional shifts across disease states (Figure 1E). Epithelial cells showed marked depletion during active disease followed by partial recovery during MH (Kruskal-Wallis test, p=0.0226). Fibroblasts were also depleted during active disease and recovered during MH (Kruskal-Wallis test, p=0.0326).

Subclustering of the epithelial compartment revealed vast heterogeneity in epithelial subset composition across disease states (Supplementary Figure 2). While the epithelial compartment overall was depleted during active disease and partially recovered during MH, some epithelial subsets exhibited more pronounced compositional shifts. Mature enterocytes (expressing *TMIGD1, AQP8*), absorptive cells (*CNNM2*, *CMBL*), and a small *BEST4/CA7^+^* epithelial population were markedly depleted in active UC and restored in MH (Supplementary Figure 2C; Kruskal-Wallis test, p=0.02, p=0.023, p=0.01, respectively, and Supplementary Table 3).

Conversely, immune populations, including plasma cells, T cells, and monocytes, were enriched during active disease (Kruskal-Wallis test, p=0.0295, p=0.005, p=0.018, respectively; Figure 1E) with plasma cells and monocytes remaining relatively elevated during MH (Figure 1E, Supplementary Figure 3A-D and Supplementary Table 4).

Beyond changes in cellular composition, both the immune compartment and the epithelial compartments exhibited persistent transcriptional alterations during MH. The immune compartment exhibited a distinct transcriptomic profile compared to healthy controls, characterized by upregulation of humoral and myeloid-associated genes (*IGHG1*, *LYZ*, *C1QB*) and downregulation of metabolic genes (*MT-CO1*, *SLC25A3*, Supplementary Figure 3E). Pathway analysis of MH-upregulated genes further revealed enrichment of phagocytosis and microbial response pathways (Supplementary Figure 3F). Similarly, despite partial restoration of epithelial subset composition, MH epithelial cells remained transcriptionally distinct from healthy controls, exhibiting increased expression of antimicrobial and immune-associated genes (LCN2, CXCL1, S100A9) together with enrichment of antimicrobial defense and immune activation pathways (Supplementary Figure 2D-E). Together, these findings indicate that despite macroscopic healing, both the colonic epithelium and immune compartments remain transcriptionally altered.

To determine whether these compositional and transcriptional alterations were accompanied by changes in tissue-wide communication networks, we next examined cell-cell interactions using CellChat ^37^, a computational tool for inferring intercellular communication networks from scRNA-seq data. Based on expression of ligands and receptors, each cell type was assigned a sender score (Figure 1F) and a receiver score (Supplementary Figure 4 and Supplementary Table 5). Overall interaction activity varied significantly across disease progression (Kruskal-Wallis test: all cell types p=0.034). Active disease was associated with increased interaction scores across most cell types, whereas MH was accompanied by a reduction in interactions to levels similar or lower than those observed in healthy control. Across all disease states, fibroblasts consistently emerged as the dominant interacting population, exhibiting significantly higher sender interaction strength than all other cell types (healthy: p= 0.0008, Active UC: p= 0.0002, MH: p = 0.0014; Wilcoxon rank-sum test, Figure 1F).

Given the prominent alterations observed in the fibroblast compartment, we next examined fibroblast-specific transcriptional changes during MH relative to healthy tissue. MH fibroblasts exhibited significant upregulation of key inflammatory and remodeling genes, including CCL11, CXCL6, and CHI3L1 (Figure 1G). Pathway analysis of MH-upregulated genes revealed enrichment of extracellular matrix organization, epithelial proliferation, and inflammatory regulation pathways (Figure 1H). These findings support the notion that MH represents an altered biological state, and implicates stromal cells, particularly fibroblasts, in its maintenance.

Repeated cycles of inflammation and healing in UC are known to cause extensive crypt damage and loss of normal tissue architecture. To examine whether the alterations in cellular compositions and cell-cell interactions observed by scRNA-seq were accompanied by structural alterations, we performed second harmonic generation (SHG) imaging followed by image analysis to visualize and quantify fibrillar collagen organization. SHG imaging of mucosal biopsies from 25 patients revealed vast alterations in collagen structure and crypt architecture in the course of UC (Figure 1I). In particular, crypt numbers were significantly reduced during active disease and failed to recover during MH (Figure 1J). The persistence of structural damage together with the elevated fibroblast signaling activity suggests that altered epithelial-stromal interactions contribute to the biologically distinct state of MH. Moreover, these findings indicate that damage incurred during active inflammation bears long term consequences for tissue integrity, even in patients who appear endoscopically healed.

### Fibroblast and pericyte subsets are spatially organized and follow distinct trajectories during UC

The alterations in stromal cell abundance and interactions, as well as the vast ECM remodeling observed, led us to perform a focused analysis of the stromal compartment, encompassing fibroblasts and pericytes. Reanalysis of 7716 cells from the clusters annotated as fibroblasts and pericytes in our scRNA-seq dataset resolved the stroma into 6 fibroblasts and 2 pericyte populations consistently observed across patients and disease states (Figure 2A-B and Supplementary Figure 5). Cluster annotation was performed using the U-method, which identified unique marker signatures for each stromal subset (Figure 2C and Supplementary Table 2). To further characterize these populations we performed differential expression and pathway enrichment analyses, revealing distinct functional programs associated with each subset (Supplementary Table 6). One fibroblast population expressing *VSTM2A* was enriched for genes associated with the upper crypt region, mainly BMP signaling, and was therefore termed “Top crypt fibroblasts” ^25^ (Figure 2C-D and Supplementary Table 6). This cluster also included canonical telocyte markers such as *FOXL1* and *PDGFRA^38,39^* (Supplementary Table 2). Conversely, a cluster of OGN⁺ fibroblasts (Figure 2C) was enriched for WNT ligands (*WNT2, WNT2B, RSPO3*) and BMP-antagonists (*GREM1*) known to support the intestinal stem-cell niche ^25^ (Figure 2D) and was therefore termed “Stem-cell-niche (SCN) fibroblasts”. A large population which we termed *“*General fibroblasts” expressed interstitial matrix genes (*FGFR2, ADAM28, APOE*); “Myofibroblasts” expressed canonical contractile markers (*HHIP, SOSTDC1,* Myofibroblasts, Figure 2C); and a cluster lacking unique markers but enriched for stress-related genes was termed “Stress fibroblasts”, possibly reflecting an artifact of the single-cell dissociation process (Figure 2C and Supplementary Table 6). An additional fibroblast cluster was characterized by high expression of chemokines (*CCL19, CCL21;* Figure 2C), resembling a subset previously shown to localize adjacent to Gut-Associated Lymphoid Tissue (GALT) and to orchestrate immune cell recruitment, and we therefore termed it “GALT fibroblasts” ^7,40^.

**Figure 2.**
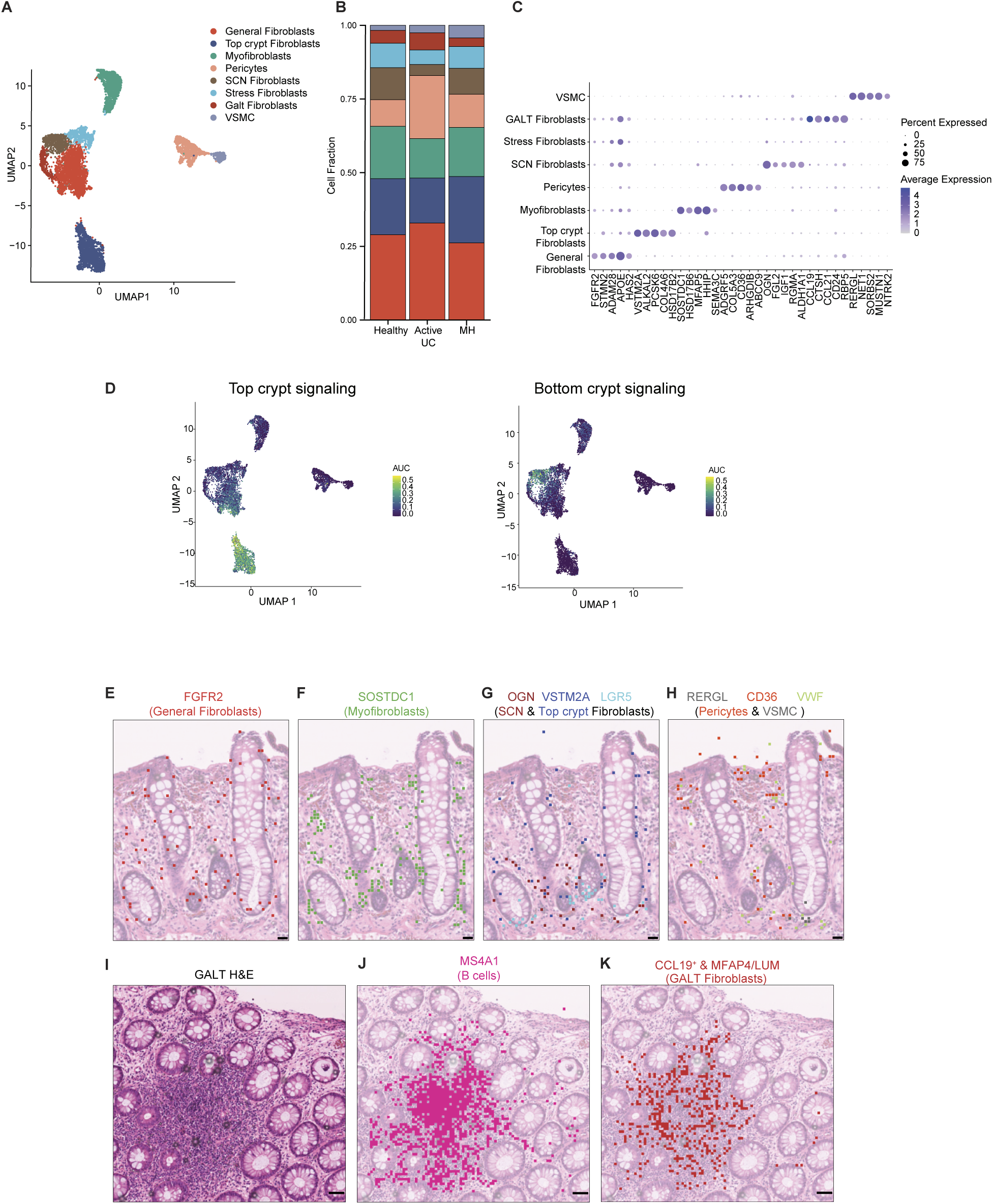
Single-cell and spatial transcriptomic analyses identify distinct fibroblast and pericyte subsets that change across UC states. (**A**) Subclustering of fibroblasts and pericytes using Seurat and UMAP visualization of the scRNA-seq dataset. **(B)** Stacked bar plot illustrating the relative proportions of the high-resolution stromal subpopulations across disease states **(C)** Dot plot displaying the top five UEGs for each of the high-resolution subclusters. **(C)** UMAP visualization of Top crypt (left; *PDGFRA, SOX6, BMP2, BMP3, BMP4, BMP5, BMP7, WNT5A, NRG1*) and Bottom crypt (right; *GREM1, RSPO3, IGF1, WNT2, WNT2B, VEGFD*) signaling activity. Color reflects the AUC enrichment. **(E–K)** Visium HD spatial transcriptomics was performed on 12 biopsies (n = 4 per disease state: healthy, active UC, and MH) at 8 µm resolution**. (E-H)** Representative spatial plots are shown from an ROI containing longitudinally oriented crypts from a patient in MH, displaying all positive spots for top UEGs overlaid on H&E sections: **(E)** General Fibroblasts (*FGFR2*); **(F)** Myofibroblasts (*SOSTDC1*); **(G)** Top crypt Fibroblasts & SCN Fibroblasts (*VSTM2A, OGN*) and LGR5 to delineate the stem cell niche; and **(H)** Pericytes and VSMCs (*CD36, RERGL*) with *VWF* included to mark the endothelium. Scale bar - 25µm **(I-K)** GALT area H&E staining **(I)** and Visium HD spatial plots **(J-K)** of a representative GALT region from a patient in MH. **(J)** Spatial distribution of the top B cell UEG (*MS4A1)* to mark the GALT area. **(K)** Spatial mapping of GALT fibroblasts, indicated by spots double-positive for the top UEG *CCL19* and for either *MFAP4 or Lum* (top fibroblast UEGs). Scale bar - 50µm.

Pericytes diverged into two distinct subsets - a small subset expressing vascular smooth muscle cell (VSMC) genes (*RERGL, MUSTN1*; Figure 2C and Supplementary Table 6) which we termed “VSMC” and a larger subset (“Pericytes”) enriched for genes involved in matrix remodeling and blood vessel development (*ADGRF5, COL5A3*; Figure 2C and Supplementary Table 6) resembling matrix-remodeling pericytes previously described by us in cancer models outside the colon ^41^.

Stromal subset compositions varied extensively across disease states (Figure 2B and Supplementary Figure 5). General fibroblasts and pericytes expanded during active disease whereas SCN fibroblasts were significantly depleted and failed to fully recover during MH(Kruskal-Wallis test, p= 0.014). These findings suggest that stromal populations follow distinct trajectories during UC progression and MH.

The divergent trajectories of stromal subsets across disease states prompt us to examine their spatial organization, *in situ*. Because the colon is a highly polarized organ and fibroblasts are known to occupy distinct positions along the crypt axis ^25^, we performed VisiumHD spatial transcriptomics on a subset of 12 patients from our cohort (Figure 1A and Supplementary Figure 6; see Methods). This analysis confirmed the identities inferred from scRNA-seq, revealed new marker genes and uncovered spatial relationships between stromal, epithelial, and immune populations. *FGFR2*^+^ general fibroblasts and *SOSTDC1*^+^ myofibroblasts ^42^ were distributed along the length of the crypt axis (Figure 2E-F). In contrast, *VSTM2A*^+^ top-crypt fibroblasts were excluded from the crypt base and segregated from *LGR5*^+^ intestinal stem cells (Figure 2G). SCN fibroblasts, for which we identified *OGN* as a unique marker, localized to the crypt base and colocalized with *LGR5*^+^ cells (Figure 2G). The two pericyte subsets were also spatially segregated. *RERGL^+^* VSMCs colocalized with *VWF^+^*endothelial cells at the bottom of the crypt, while the larger *CD36^+^* pericyte population was distributed throughout the mucosa and only partially colocalized with endothelial cells (Figure 2H). Finally, *CCL19*^+^ GALT fibroblasts, also marked by *MFAP4* or *LUM* which we identified as fibroblast top UEGs (Figure 1D) colocalized with *MS4A1*^+^ B cells within GALT, supporting their designation as GALT fibroblasts (Figure 2I-K). Together, these findings establish a spatially organized stromal landscape that undergoes extensive remodeling during UC progression and mucosal healing.

### Inflammatory fibroblasts emerge during active disease while stem-cell-niche fibroblasts diminish and remain altered in MH

The divergent trajectories observed within the stromal compartment prompted us to examine transcriptional heterogeneity among fibroblasts in greater detail. We first asked whether the expansion of the General Fibroblast population during active disease could be driven by the emergence of a distinct inflammatory subpopulation masked within the General Fibroblast cluster. Reanalysis of General Fibroblasts using Seurat clustering and UMAP visualization revealed a clear segregation of cells derived from patients with active UC compared with those from controls (Figure 3A and Supplementary Figure 7A-B). Based on their relative enrichment and transcriptional profiles, we defined the subpopulation predominantly enriched in healthy tissues as “Normal fibroblasts”, representing a homeostatic fibroblast signature, whereas the subset expanded during active inflammation was designated as “Inflammatory fibroblasts” (Supplementary Table 2). In active UC, inflammatory fibroblasts dominated the general fibroblast compartment, accounting for more than 75% of cells within this population (Supplementary Figure 7B). In contrast, MH exhibited an intermediate and highly variable phenotype, with inflammatory fibroblasts comprising 15-60% of the general fibroblast population (Supplementary Figure 7B).

**Figure 3.**
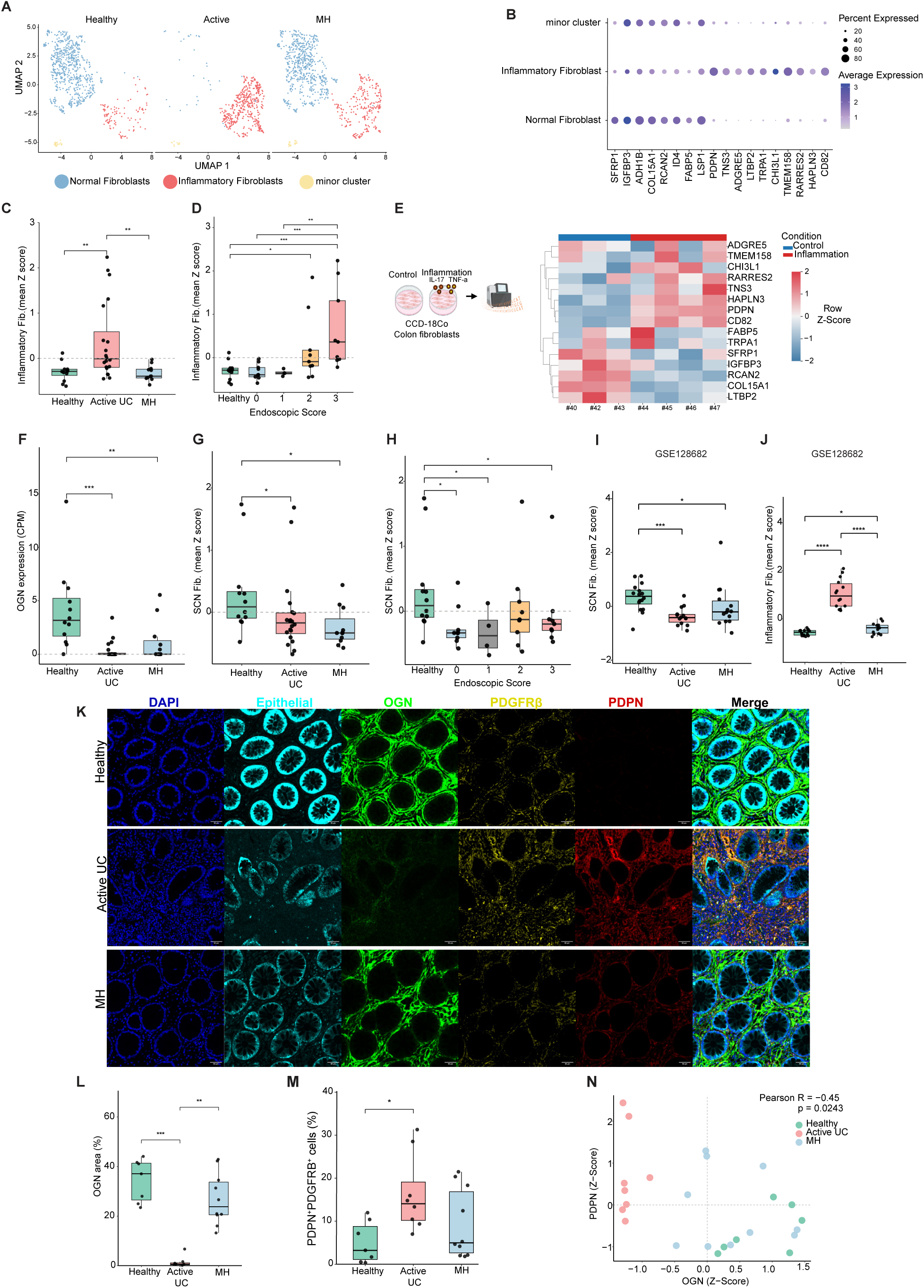
Single-cell and bulk RNA-seq reveal expansion of inflammatory fibroblasts and depletion of SCN fibroblasts across UC disease states. **(A)** Subclustering of the General Fibroblast population from the scRNA-seq dataset using Seurat and UMAP visualization, split by disease state. **(B)** Dot plot showing the top UEGs defining Normal and Inflammatory fibroblast subsets, identified using the U-method algorithm **(C-D)** Bulk RNA-seq analysis of 43 colonic biopsies spanning all disease states (n = 12 healthy, 20 active UC, and 11 MH). The Inflammatory Fibroblast signature is shown as boxplots of mean z-scores across disease states **(C)** and stratified by endoscopic score, ranging from 0 (endoscopic remission, MH) to 3 (most severe disease activity) **(D). (E)** Schematic overview of the *in vitro* experimental design (left) and heatmap of bulk RNA-seq data (right) from primary human colonic fibroblasts (CCD-18Co) cultured with or without inflammatory cytokines (IL-17 and TNFα) for 24 hours. The heatmap shows expression of UEGs defining the Normal and Inflammatory fibroblast signatures (as in C) that are detected in the bulk RNA-seq data. **(F)** Boxplots showing bulk RNA-seq expression of the SCN marker *OGN* across disease states in the cohort analyzed in **(C)**. **(G-H)** Boxplots showing mean z-scores of the SCN fibroblast signature across disease states **(G)** and stratified by endoscopic score **(H)** in the bulk RNA-seq cohort analyzed in **(C)**. **(I-J)** Boxplots showing mean z-scores of the SCN fibroblast **(I)** and Inflammatory fibroblast **(J)** signatures in an independent, publicly available bulk RNA-seq dataset (GSE128682) across disease states. Statistical significance was determined using pairwise Wilcoxon rank-sum tests (*p < 0.05, **p < 0.01, ***p < 0.001). **(K-N)** Multiplexed immunofluorescence (MxIF) staining of colonic sections across UC stages (n=7 healthy, 8 active UC and 10 MH), using antibodies against epithelial cells (a cocktail of CK18, CK7, EpCAM; cyan), PDGFRβ (pan-fibroblast; yellow), OGN (SCN fibroblasts; green), and PDPN (inflammatory fibroblasts; red). DAPI was used to mark nuclei (blue). Representative images are shown in **(K).** Scale bar - 50 µm. **(L-N)** Quantification of MxIF images across disease states. **(L)** Pixel-based quantification showing the percentage of stromal area positive for OGN. **(M)** Cell-based quantification showing the fraction of PDGFRβ^+^ cells expressing PDPN. Group comparisons were assessed by Kruskal-Wallis test with Dunn’s post-hoc test, Benjamini-Hochberg adjusted (*p < 0.05, **p < 0.01, ***p < 0.001). **(N)** Pearson correlation analysis of normalized OGN and PDPN protein levels across individual patients, based on the MxIF staining.

This inflammatory fibroblast subset was characterized by upregulation of *CHI3L1, PDPN*, and *RARRES2* (Figure 3B), markers associated with inflammatory cancer-associated fibroblasts (iCAFs) in the tumor microenvironment^28,43,44^. Indeed, pathway enrichment analysis using Metascape ^45-47^ revealed suppression of developmental programs and enrichment of ECM remodeling and inflammatory pathways, suggesting convergence between stromal remodeling programs in chronic inflammation and cancer (Supplementary Table 7).

To validate the enrichment of inflammatory stromal programs in active UC, we analyzed our bulk RNA-seq dataset derived from a cohort of 43 patients spanning all disease stages (Figure 1A). An inflammatory signature composed of the top 10 UEGs identified by scRNA-seq was significantly enriched in active UC compared to healthy controls and patients in MH (Figure 3C and Supplementary Table 8; see methods). Given the clinical heterogeneity of UC, we next examined whether this inflammatory signature varied with disease severity. Stratification of patients by endoscopic disease activity (Mayo endoscopic score; MES) showed that the inflammatory signature increased progressively with disease severity, with highest scores in patients with severe disease (MES=3; Figure 3D). Furthermore, stratification of the cohort by age revealed that the disease-associated trends in stromal signatures were similar in both pediatric and adult patients (Supplementary Figure 7C).

To test whether this transcriptional program could be directly induced by inflammatory cues on normal fibroblasts, we cultured CCD-18Co normal human colon fibroblasts in the presence of IL-17 and TNFalpha ^48,49^ for 24h after which we performed bulk RNA-seq. We then queried the expression of the top UEGs defining the normal and inflammatory fibroblast signatures (Figure 3C). Of the top UEGs defining the normal and inflammatory fibroblast signatures (Figure 3E). Of those genes detected in the CCD-18Co cells, cytokine stimulation induced a clear transcriptional shift, with downregulation of normal fibroblast markers and upregulation of inflammatory fibroblast markers, particularly *PDPN, HAPLN3* and *CD82,* indicating that inflammatory cues not only induce inflammatory fibroblast programs but are accompanied by suppression of normal fibroblast features (Figure 3E).

Concomitant with the emergence of inflammatory fibroblasts, our scRNA-seq analysis revealed a marked depletion of SCN fibroblasts during active UC, with incomplete recovery during MH (Figure 2B). We also identified OGN as a novel, highly specific marker for the SCN fibroblast subset, both within the stromal compartment and across the full scRNA-seq dataset (Figure 2C and Supplementary Figure 7D). To further interrogate this subset, we examined OGN expression in our bulk RNA-seq cohort. OGN expression was significantly reduced in active UC and remained significantly lower than in healthy controls during MH (Figure 3F). This effect was not restricted to OGN alone. Analysis of a signature composed of the top 10 UEGs defining the SCN population confirmed that this subset is depleted in active UC and remains significantly depleted also in MH (Figure 3G-H and Supplementary Table 2; see also similar trends across pediatric and adult patients in Supplementary Figure 7E).

To assess the generalizability of our findings, we evaluated both the inflammatory and SCN fibroblast signatures in an independent, publicly available bulk RNA-seq dataset comprising 44 patients across disease stages (n=16 healthy controls, n=14 active UC and n=14 remission (MH, see methods); GSE128682)^22^. Consistent with our findings, the external dataset confirmed enrichment of the inflammatory fibroblast signature in active UC and depletion of the SCN signature across disease stages (Figure 3I-J). Notably, in this cohort the inflammatory signature remained significantly elevated in MH compared to healthy controls, indicating variability in the extent of inflammatory resolution. Together, these findings indicate that MH is defined by robust depletion of SCN-supporting fibroblasts and variable persistence of inflammatory fibroblast programs, suggesting that impaired epithelial niche support and residual inflammation together underlie incomplete tissue restoration.

### Spatial and protein-level analyses reveal persistent imbalance between inflammatory and niche-supporting fibroblasts during UC progression

To examine how the transcriptional transitions in fibroblast states are manifested in tissue architecture, we analyzed their spatial distribution *in situ*. We focused on the top UEGs defining each population: OGN for the SCN fibroblasts, and PDPN, a well-established marker of CAFs that has also been implicated as a marker of IBD^7,8,26,28,50^, for the inflammatory subset. Visium HD analysis showed increased PDPN expression during active disease, with partial reduction during MH (Supplementary Figure 7F). In contrast, OGN was abundant near LGR5^+^ epithelial cells in the stem cell niche in healthy mucosa but markedly reduced in both active UC and MH, indicating persistent loss of SCN-supporting fibroblasts despite clinical remission (Supplementary Figure 7G).

We next sought to robustly quantify these dynamics at the protein level. To achieve this, we performed multiplexed immunofluorescence (MxIF) staining for PDPN and OGN. PDGFRβ was used as a pan-stroma marker and the combination of CK18, CK7 and EpCAM staining delineated epithelial crypt architecture (Figure 3K). In healthy controls, OGN, a secreted protein, was robustly expressed in stromal regions adjacent to crypts, whereas PDPN expression was low. In stark contrast, chronic inflammation was associated with near-complete loss of OGN and a significant increase in PDPN^+^/PDGFRβ^+^ inflammatory fibroblasts (Figure 3L-M). During MH, OGN expression showed significant recovery and PDPN levels were reduced to intermediate levels. Correlation analysis confirmed an inverse relationship between these two markers, with high OGN/ low PDPN in healthy tissue, low OGN/high PDPN in active UC, and intermediate, variable levels during MH (Figure 3N). Together, these findings indicate that MH is characterized by a persistent imbalance between inflammatory and niche-supporting fibroblast states. Notably, the variability observed across patients suggests that MH represents a heterogeneous biological state, in which incomplete stromal recovery and retention of an intermediate fibroblast phenotype may reflect a form of stromal “memory” of prior inflammation and potentially mark tissue at risk for disease relapse.

### UC drives expansion and phenotypic reprogramming of pericytes beyond canonical vascular support

While fibroblasts constitute the dominant stromal population and undergo extensive remodeling in UC, our scRNA-seq analysis also highlighted significant compositional and signaling shifts within the pericyte compartment across disease stages (Figure 2B and Supplementary Figure 5). To further examine these changes, we first sought to validate pericyte expansion in the bulk RNA-seq cohort. Applying a pericyte-specific signature composed of the top 10 UEGs identified by our scRNA-seq analysis (Supplementary Table 8), we observed that pericytes tend to expand in active UC and decrease during MH to levels comparable to healthy controls (Figure 4A). Stratification by endoscopic disease activity supported this pattern, showing a progressive increase in pericyte signature scores with disease severity, with low levels in healthy controls and patients in MH (MES=0), and the highest scores in patients with severe disease (MES=3; Figure 4B), indicating that pericyte expansion closely tracks inflammatory burden in UC. These observations were corroborated in the external bulk RNA-seq dataset (GSE128682), in which pericyte signature scores were significantly increased in active disease and reduced in remission (MH), supporting the robustness of this pattern across independent cohorts (Figure 4C).

**Figure 4.**
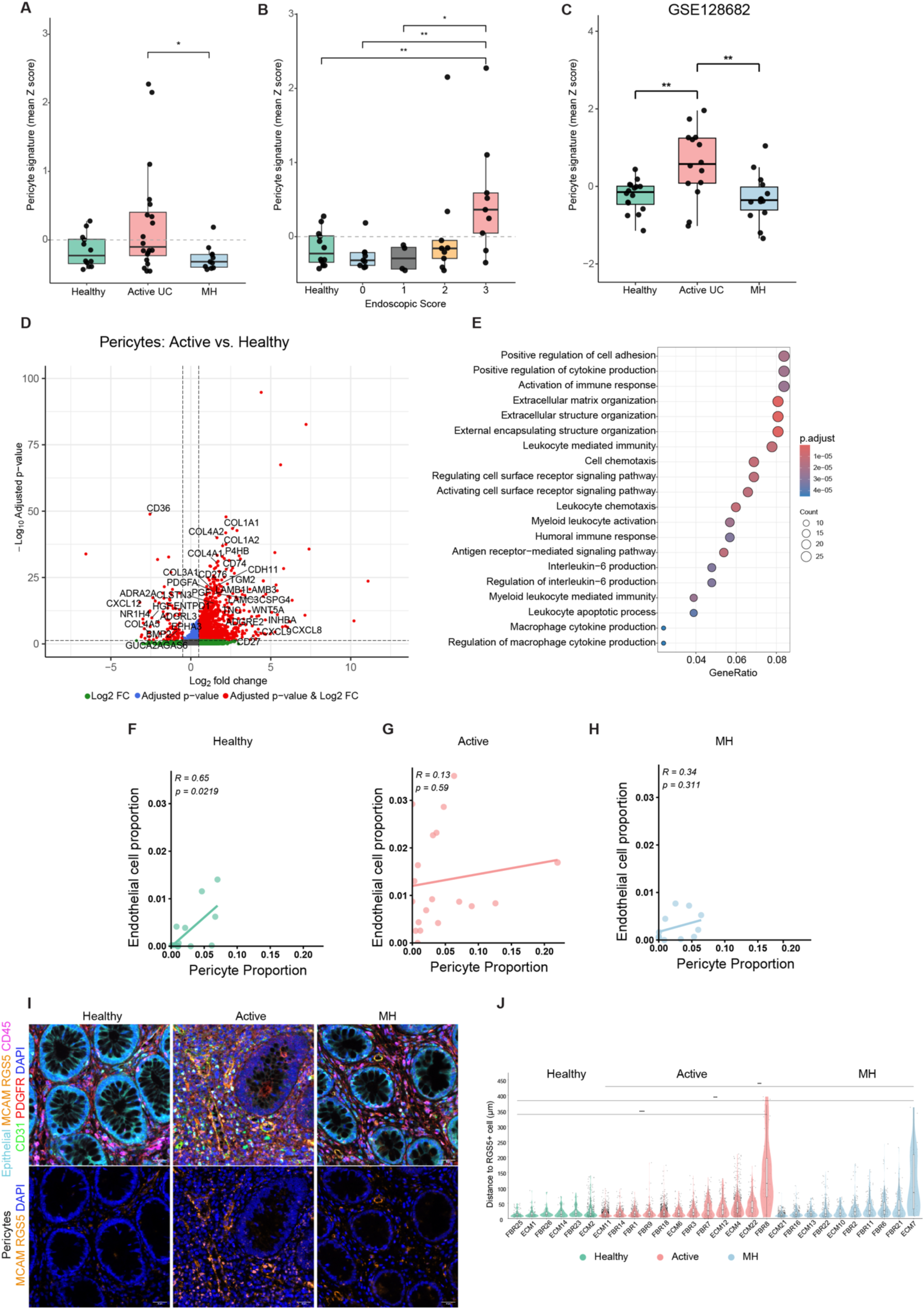
Pericytes expand and transition from vascular support to matrix-remodeling states during UC. **(A-B)** Boxplots showing mean z-scores of the pericyte signature across disease states **(A)** and stratified by endoscopic score **(B)** in the bulk RNA-seq cohort analyzed in Figure 3E. **(C)** Boxplots showing mean z-scores of the pericyte signature across disease states in an independent publicly available bulk RNA-seq dataset (GSE128682), as analyzed in Figure 3L. **(D)** Volcano plot showing differential expression of ligand- and receptor-encoding genes in pericytes from active UC and healthy controls, identified using CellChat analysis of the scRNA-seq dataset. Colors indicate significance levels based on defined thresholds: Green (|log2FC| ≥ 0.5, p.adj ≥ 0.05) blue (|log2FC| ≤ 0.5, p.adj < 0.05), red (|log2FC| ≥ 0.5, p.adj < 0.05) and grey (not significant). **(E)** Pathway enrichment analysis of differentially expressed genes in pericytes from active UC versus healthy controls using clusterProfiler. Enrichment was performed on significantly upregulated genes (log₂ fold change > 1.5, adjusted p value < 0.05) identified from the scRNA-seq data. **(F–H)** Scatter plots showing the correlation between endothelial and pericyte proportions, estimated by MuSiC deconvolution of the bulk RNA-seq data, in healthy **(F)**, active UC **(G)**, and MH **(H)** patients. Pearson correlation coefficients (R) and p-values are indicated. **(I-J)** MxIF staining of colonic sections across UC stages (n=6 healthy, 12 active UC and 10 MH), using antibodies against epithelial cells (a cocktail of CK18, CK7, EpCAM; cyan), MCAM and RGS5 (pericytes; orange), CD45 (immune cells; magenta), CD31 (endothelial cells; green), and PDGFRβ (pan-stroma; yellow). DAPI was used to mark nuclei (blue). Representative images are shown in **(I)**, with lower panels highlighting pericyte markers (MCAM, RGS5). Scale bar - 50 µm. **(J)** Violin plots showing the distribution of distances between pericytes and the vasculature, calculated as the minimal Euclidean distance (µm) from CD31^+^ endothelial cells to the nearest RGS5^+^ pericyte across individual patients in healthy, active UC, and MH groups. Statistical differences were evaluated using a two-sample Kolmogorov–Smirnov (KS) test, with Benjamini-Hochberg correction for multiple comparisons.

To assess whether pericytes also undergo transcriptional changes during inflammation, we built on our global CellChat analysis of intercellular communication across UC stages (Figure 1F) and compared the expression of ligand- and receptor-encoding genes in pericytes from active UC and healthy controls (Figure 4D). This analysis revealed broad transcriptional remodeling, characterized by downregulation of canonical pericyte markers (e.g., *CD36*) and upregulation of fibroblast-associated genes (*COL1A1, COL1A2, COL4A2, INHBA*) involved in adhesion, cytokine production and matrix remodeling, suggesting a shift toward a more mesenchymal, matrix-remodeling phenotype (Figure 4D-E).

To further characterize this transition, we performed pseudotime analysis using Monocle3 ^51^. The trajectory was rooted in cells exhibiting the highest expression of the canonical pericyte marker *RGS5* and revealed a bifurcating progression into two distinct axes (Supplementary Figure 8A-B). One branch remained relatively close to the origin, representing a shift toward a more VSMC-like state. In contrast, the second branch extended along pseudotime, reflecting more profound transcriptomic alterations. To define the molecular programs underlying this transition, we evaluated gene expression dynamics along the trajectory. Using Moran’s I statistic, we identified and clustered the most significantly dynamic genes across pseudotime (Supplementary Figure 8C and Supplementary Table 9). Early pseudotime states were enriched for pericyte associated genes (*STEAP4*, *BCAM*, *RCAN2*) and contractile markers (*MYH11*, *FLNA*, *MYL9*). In contrast, late pseudotime states showed prominent upregulation of ECM components and remodeling enzymes (*COL1A1*, *MMP2*, *LOXL2*). Notably, this transition involved a transient early upregulation of *TAGLN* and *ACTA2*, consistent with early myofibroblast activation, a pattern previously described also in liver regeneration ^52^. Collectively, these dynamics suggest that under inflammatory conditions, pericytes progressively lose their canonical identity and acquire a matrix-remodeling, fibroblast-like phenotype. Pericytes are mural cells classically implicated in angiogenesis and vascular homeostasis ^53^. Nevertheless, none of these pathways were upregulated in active UC (Figure 4E). To assess whether pericyte expansion and transcriptional rewiring were accompanied by altered coupling to endothelial cells, we examined the relationship between endothelial and pericyte fractions using MuSiC^54^ deconvolution of the bulk RNA-seq data (Figure 4F-H). In healthy tissue, endothelial and pericyte fractions were strongly correlated (R=0.65), consistent with their canonical vascular association. This relationship was markedly disrupted in active UC (R=0.13), and only partially restored during MH (R=0.34), indicating that inflammation-associated pericytes persist with functions that are increasingly independent of vascular support.

The decoupling of pericytes from endothelial cells is expected to affect their spatial localization within the tissue. To test this, we performed MxIF and quantified the distance between CD31^+^ endothelial cells and RGS5^+^ pericytes. CD45 and epithelial cell cocktail (CK18, CK7, EpCAM) were used to exclude immune cells and epithelium, respectively (Figure 4I). In healthy mucosa, pericytes were predominantly restricted to the perivascular niche. In contrast, during both active UC and MH, pericytes were frequently observed dispersed throughout the lamina propria (Figure 4I). To quantify this redistribution, we measured the distance between CD31⁺ endothelium and RGS5⁺ pericytes using Qupath^55^ image analysis and assessed changes in the distribution using a Kolmogorov–Smirnov (KS) test. This analysis revealed a significant shift in distance distributions. Specifically, in patients with active disease, pericytes were observed at substantially increased distances from the endothelium compared to healthy controls, and this phenomenon was also maintained in patients with MH (Figure 4J). These findings suggest that during UC, a subset of pericytes may undergo a mural-to-mesenchymal transition, potentially detaching from the vasculature to participate in the broader remodeling of the inflammatory stroma. This reprogrammed state partially persists during MH, suggesting a sustained contribution of pericytes to stromal remodeling and incomplete tissue restoration.

### Stromal remodeling signatures predict therapeutic response in UC

The persistent stromal remodeling observed across disease stages prompted us to examine whether these alterations are associated with clinical outcomes and therapeutic response. To this end, we analyzed a publicly available dataset (GSE16879) tracking responses in 24 patients with active UC to infliximab (IFX), an anti-TNF antibody, alongside 6 healthy controls ^56^. At baseline (pre-treatment), signatures of inflammatory fibroblasts and pericytes were significantly elevated relative to healthy controls, while the SCN fibroblast signature showed a trend toward reduction (Figure 5A-B and Supplementary Figure 8). Strikingly, both inflammatory fibroblast and pericyte signatures were significantly higher in patients who subsequently were classified as infliximab non-responders (NR) compared with responders (R) and healthy controls, indicating that baseline stromal alterations may predict therapeutic outcome (Figure 5A–B). Analysis of post-treatment samples revealed that these signatures persisted in non-responders, whereas responders showed a marked reduction, reaching levels comparable to those observed in healthy controls (Figure 5C-D).

**Figure 5.**
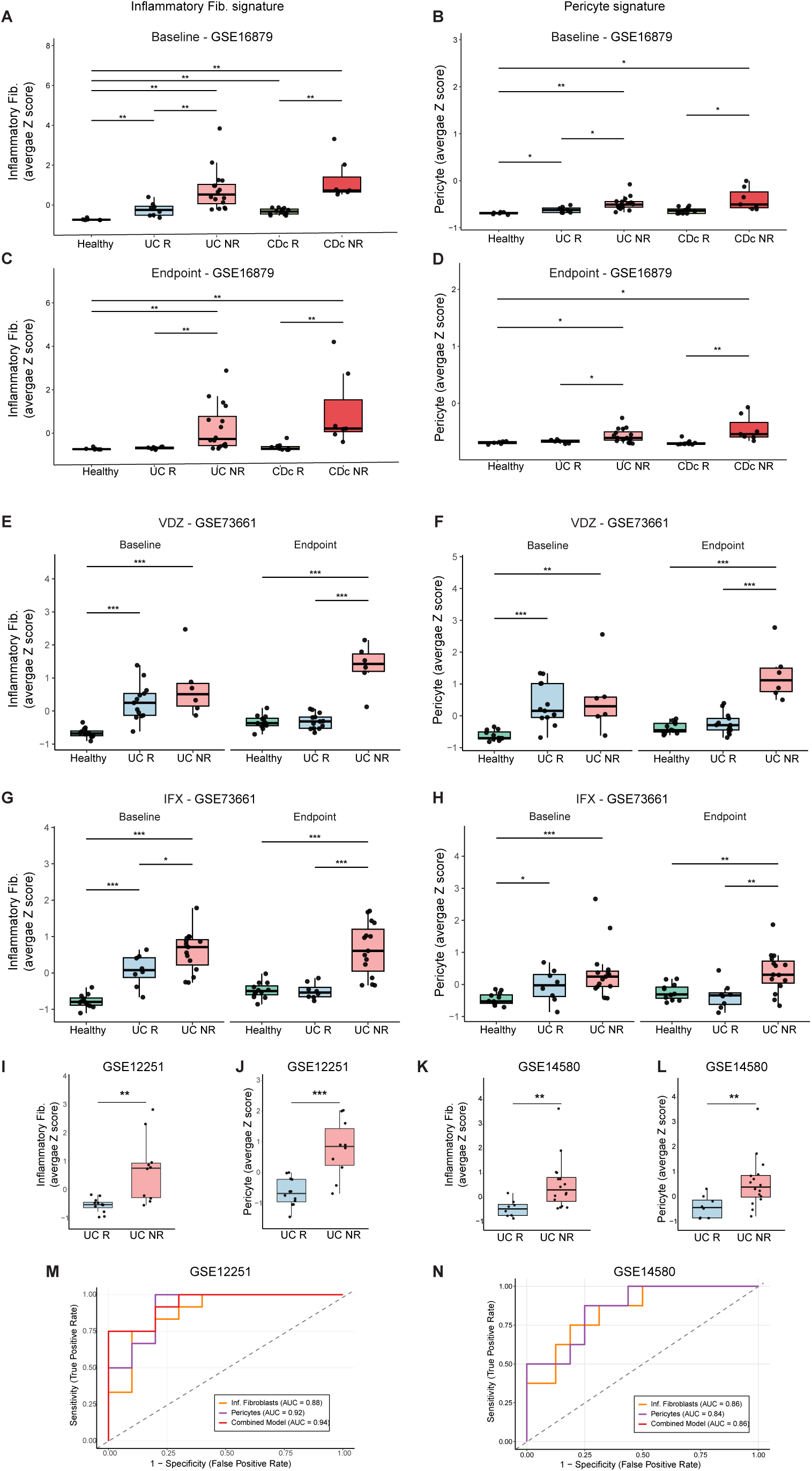
Baseline stromal signatures of inflammatory fibroblasts and pericytes predict therapeutic response to anti-TNFα therapy in UC patients. **(A-B)** Boxplots displaying baseline (pre-treatment) Z-scores of the inflammatory fibroblast **(A)** and pericyte **(B)** signatures in healthy controls (n=6), UC (n=24) and colonic Crohn’s disease (CDc; n=19) patients from a publicly available dataset (GSE16879) tracking responses to the anti-TNF therapy Infliximab (IFX). Patients are stratified by their future response to Infliximab (IFX). **(C–D)** Boxplots displaying post-treatment Z-scores of the inflammatory fibroblast **(C)** and pericyte **(D)** signatures in the same patient cohort (GSE16879). **(E–F)** Boxplots displaying Z-scores of the inflammatory fibroblast **(E)** and pericyte **(F)** signatures in an independent cohort (GSE73661) including healthy controls (n=12) and active UC patients treated with Vedolizumab (VDZ, n=23), evaluated at baseline and at Week 52 (endpoint). **(G–H)** Boxplots displaying Z-scores of the inflammatory fibroblast **(G)** and pericyte **(H)** signatures in the same cohort (GSE73661) for patients treated with IFX (n=19), evaluated at baseline and post-treatment endpoint. **(I–N)** Evaluation of baseline stromal signatures as predictors of IFX response in two external UC cohorts: GSE12251 (n=22 patients) and GSE14580 (n=24 patients). Boxplots display baseline Z-scores of the inflammatory fibroblast **(I, K)** and pericyte **(J, L)** signatures, stratified by future therapeutic outcome. Receiver operating characteristic (ROC) curves **(M-N)** evaluate the predictive performance of the individual stromal signatures and a combined logistic regression model in each cohort. Statistical significance across boxplots was assessed using Wilcoxon rank-sum tests (*p < 0.05, **p < 0.01, ***p < 0.001).

The dataset also included 19 patients with colonic Crohn’s disease (CDc). Analysis of stromal signatures in this cohort revealed similar patterns to those observed in the UC cohort - elevated levels of inflammatory fibroblast and pericyte signatures in patients with active disease, which failed to recover in the non-responders, post-treatment with IFX (Figure 5A-D). SCN fibroblast signatures showed variable patterns, with reduced levels in responders, but values comparable to healthy controls in non-responders, suggesting potential disease-specific differences in SCN dynamics (Supplementary Figure 9).

To further evaluate stromal signatures across therapeutic modalities, we analyzed an independent dataset ^57^ (GSE73661) including RNA microarray data from UC patients treated with either the anti-α4β7 intgerin antibody vedolizumab (VDZ, n=19 patients; Figure 5E-F), or IFX (n=23 patients; Figure 5G-H). For the VDZ cohort, clinical remission at Week 52 was chosen as the endpoint (see methods). The SCN fibroblast signature was excluded from this analysis due to incomplete representation of its marker genes in this dataset. Consistent with our earlier observations, pre-treatment samples showed elevated inflammatory fibroblast (Figure 5E, 5G) and pericyte signatures (Figure 5F, 5H) in both future responders and non-responders compared to healthy controls. Following treatment, responders showed a significant reduction in these signatures, with the pericyte signature normalizing to levels comparable to healthy controls. In contrast, non-responders maintained significantly elevated levels of both inflammatory fibroblast and pericyte signatures (Figure 5E-H).

To further evaluate the predictive potential of baseline pericyte and inflammatory fibroblast signatures, we analyzed two additional independent datasets (GSE12251 and GSE14580)^56^ comprising RNA microarray data from patients prior to IFX treatment, with known therapeutic outcomes (n = 12 responders and 10 non-responders in GSE12251; n = 8 responders and 16 non-responders in GSE14580). Consistent with our previous analyses, non-responders in both cohorts exhibited significantly higher baseline enrichment of inflammatory fibroblast and pericyte signatures compared to future responders (Figure 5I-L). Receiver operating characteristic (ROC) analysis confirmed the predictive power of these baseline stromal signatures (Figure 5M-N). In the GSE12251 dataset, this yielded high Area Under the Curve (AUC) values for inflammatory fibroblasts (AUC = 0.88), pericytes (AUC = 0.92), and the combined model (AUC = 0.94) (Figure 5M). This predictive capacity was replicated in the GSE14580 dataset, which demonstrated AUC values of 0.86 for inflammatory fibroblasts, 0.84 for pericytes, and 0.86 for the combined model (Figure 5N).

Together, these findings demonstrate that stromal remodeling is strongly associated with therapeutic outcomes. Inflammatory fibroblast and pericyte signatures are predictive of treatment response at baseline, while successful therapy shifts the stromal state toward a MH-like configuration rather than full restoration of healthy tissue architecture.

### Spatial transcriptomic analysis reveals extensive domain-specific remodeling of the colonic mucosa during UC and MH

The marked remodeling of fibroblast and pericyte populations, and the association with clinical response, prompted us to ask how stromal reprogramming reshapes the architectural organization of the colonic mucosa. We therefore turned to our Visium HD spatial transcriptomics data and performed spatial domain analysis using KNN clustering via the 10x Loupe Browser to define transcriptionally coherent anatomical regions across disease states. This unsupervised approach consistently partitioned the tissue into four principal compartments observed across controls, UC patients with active disease and UC patients exhibiting MH states: the crypts, the luminal-proximal top crypts, the lamina propria (LP), and gut-associated lymphoid tissue (GALT; Figure 6A, Supplementary Figure 10A-C and Supplementary Table 10). A fifth compartment corresponding to the muscularis mucosa was detected in healthy and MH states.

**Figure 6.**
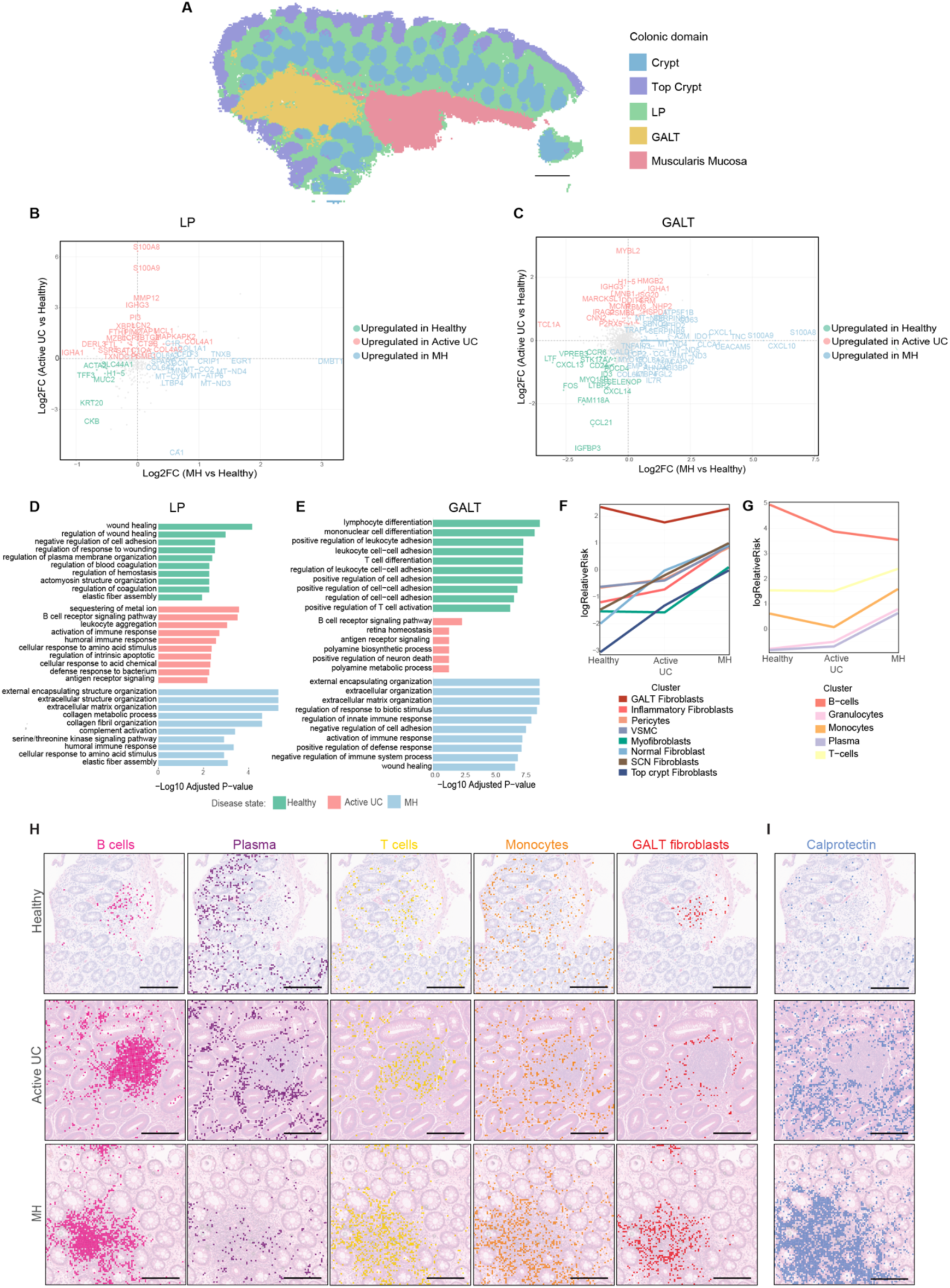
Spatial transcriptomics reveals compartment-specific remodeling of the mucosa that fails to resolve during MH. **(A)** Visium HD spatial transcriptomics data were analyzed using KNN clustering to define tissue compartments across disease states. A representative spatial map of healthy control tissue (see Supplementary Figure 5A for corresponding H&E) is shown, colored by domain: Crypts (blue), Top Crypts (purple), Lamina Propria (LP, green), GALT (yellow), and Muscularis Mucosa (red). Scale bar - 200µm **(B-C)** Scatter plots showing domain-specific gene expression changes within the LP **(B)** and GALT **(C)**. Genes were categorized based on differential expression (|log2FC| > 0.5) and colored according to the disease state in which they are upregulated: healthy (green), active UC (red) and MH (blue). **(D-E)** Bar plots showing the top 10 enriched pathways in the LP **(D)** and GALT **(E)** across disease states: healthy (green), active UC (red), and MH (blue). Pathways were identified using clusterProfiler. **(F-G)** Area enrichment analysis of stromal **(F)** and immune **(G)** cell subsets within the GALT domain. Plots show the Log relative risk (log RR) for each cell type across disease states. **(H)** Visium HD spatial maps focusing on GALT regions across disease states (healthy, active UC, and MH). Each column displays the spatial localization of specific cell subsets, identified by expression of the top UEGs from the single cell analysis: MS4A1 (B cells), MZB1 (Plasma cells), CD3E (T cells), and AIF1 (Monocytes). GALT fibroblasts are identified by spots showing co-expression of CCL19 with either MFAP4 or LUM. **(I)** Visium HD spatial mapping of the inflammatory biomarker calprotectin across disease states. spots indicate spatial regions expressing S100A8 and/or S100A9. Scale bar - 200µm

We next examined how the transcriptional composition of these spatial domains changed across disease states (Figure 6B-C and Supplementary Figure 10 D-F). Key inflammatory response genes, such as *S100A9* and *LCN2*, were upregulated in multiple domains during active UC. Beyond these global inflammatory signals, spatial analysis revealed extensive domain-specific transcriptional remodeling. For example, in the crypt area the epithelial oxidative stress factor *DUOX2* and its maturation factor *DUOXA2* were robustly upregulated in active UC (Supplementary Figure 10 D-E). Inflammatory genes involved in innate immune cell recruitment and responses such as *TNIP3, TGM2, NOS2, CXCL1* were upregulated in the top crypt region (Supplementary Figure 10 E), and *MYBL2*, a cell cycle regulator known to be expressed in CD4+ T cells ^58^, was upregulated in the GALT (Figure 6C).

MH displayed a distinct transcriptional program. Across domains, we observed upregulation of mitochondrial genes, including *MT-ND3* and *MT-ND4*, suggesting long-lasting processes of metabolic restoration (Figure 6B-C). Intriguingly, the stroma-enriched LP and GALT regions in MH samples exhibited persistent "transcriptomic scars" of prior inflammation; we identified a significant upregulation of the ECM glycoprotein encoding genes *TNC* in the GALT and *TNXB* in the LP of MH patients, compared to healthy controls, indicating persistent stromal remodeling (Figure 6B-C). Functional pathway analysis further supported this transition. Whereas the healthy LP was enriched for wound healing and adhesion pathways supporting crypt-niche maintenance, active UC showed a pronounced shift toward immunological activation and immune cell recruitment. During MH, however, the LP signature was characterized by ECM organization and collagen fibril assembly pathways (Figure 6D and Supplementary Table 11).

### The GALT adopts a distinct inflammatory configuration during MH

Domain-specific analysis of the GALT revealed pronounced shifts in stromal and immune pathways (Figure 6E). Healthy tissue was enriched for programs associated with steady-state immune surveillance and homeostasis, characterized by robust lymphocytes differentiation and regulation of leukocyte adhesion. In contrast, active UC was characterized by enrichment of humoral immune responses and metabolic stress pathways, including B cell receptor signaling and polyamine biosynthesis, consistent with the marked plasmacytosis^59,60^ and immune cell proliferation observed in active disease. During MH, the GALT exhibited a distinct transcriptional profile, enriched for innate immune regulation and ECM remodeling, indicating persistent functional reprogramming following inflammation (Figure 6E). To understand whether this transcriptional shift was driven by changes in cellular composition, we performed an area enrichment analysis comparing the GALT compartment with all other spatial domains defined by KNN clustering. Using cell-type-specific U-marker signatures to identify cell-type-enriched spots, we calculated the log relative risk (RR) of each cell type within GALT relative to other tissue domains across disease states (Supplementary Table 11). Positive log RR values indicate enrichment within GALT, whereas negative values reflect relative depletion (Figure 6F-G). In healthy controls, B cells, T cells, monocytes and CCL19+ GALT fibroblasts were enriched within the GALT. During active UC, this composition shifted markedly, with strong depletion of B cells, monocytes, and GALT fibroblasts from the GALT compartment. In MH, monocytes and GALT fibroblasts repopulated the GALT, whereas B cells were further depleted. In parallel, T cells were strongly enriched, and additional immune populations not observed in healthy or active UC GALTs, including plasma cells and granulocytes, populated the compartment together with multiple stromal subsets (Figure 6H). These changes suggest a redistribution of lymphocyte populations during disease progression, potentially reflecting migration of B cells out of GALTs and differentiation into plasma cells, alongside expansion or recruitment of T cells during MH. Curiously, S100A9 and S100A8, which together form the calprotectin complex and are hallmark markers of active inflammation, were markedly overexpressed in the crypt and lamina propria during active disease (Figure 6B and Supplementary Figure 10D) but showed selective enrichment within the GALT during MH (Figure 6C and 6I). Together with the remodeling of granulocyte and monocyte populations in this compartment, this pattern suggests that the GALT adopts a distinct inflammatory configuration during MH that is not observed in either healthy tissue or active disease, potentially reflecting a locally primed immune niche that may influence future inflammatory responses despite apparent clinical resolution.

## Discussion

Chronic inflammatory diseases such as UC are characterized by cycles of relapse and remission, yet the tissue states underlying disease progression and recovery remain poorly understood. While clinical management aims to achieve MH, high relapse rates indicate that resolution of visible inflammation does not necessarily reflect restoration of biological homeostasis. A central challenge in disease management is therefore to understand how tissue architecture and cellular programs evolve across disease states and contribute to disease persistence and recovery. Stromal cells, and fibroblasts in particular, have emerged as key regulators of tissue organization and immune responses; however, their role in shaping reslolution of the inflammatory process, and then disease recurrence in UC, remains incompletely defined. Here, we address this gap by integrating single-cell, bulk, and spatial transcriptomic analyses to define stromal cell states across UC disease stages. We identify three major stromal alterations that characterize UC progression and healing: emergence of inflammatory fibroblasts, depletion of SCN fibroblasts, and expansion of pericytes that acquire matrix-remodeling features.

Fibroblasts are increasingly recognized as heterogeneous and plastic cells capable of adopting distinct functional states in response to environmental cues. In cancer and chronic inflammatory conditions, fibroblast populations have been shown to transition into inflammatory and matrix-remodeling phenotypes, including inflammatory cancer-associated fibroblasts (iCAFs), which regulate immune responses and tissue remodeling. Our findings extend this framework to UC, demonstrating that inflammatory fibroblasts emerge from the general fibroblast pool during active disease and persist during MH. Mechanistically, we show that this pathogenic fibroblast state can be induced from normal human colonic fibroblasts by inflammatory cytokines (IL-17 and TNFα), supporting the notion that inflammatory fibroblasts arise from resident mucosal fibroblasts in response to inflammatory cues. We also observed marked inter-patient variability in the persistence of these inflammatory fibroblasts during MH, suggesting that incomplete resolution of fibroblast activation may underlie heterogeneity in disease course.

The transcriptional profile of inflammatory fibroblasts closely parallels that of iCAFs, raising important clinical implications: Patients who retain a higher burden of this inflammatory fibroblast state during clinical remission may be at increased risk of disease relapse. Moreover, the sustained presence of an iCAF-like stromal program could promote a pro-tumorigenic microenvironment, providing a potential mechanistic link to the increased risk of colitis-associated colorectal cancer (CAC) in UC patients.

In parallel with the emergence of inflammatory fibroblasts, we observed a marked depletion of SCN fibroblasts, a population defined by expression of key Wnt-supporting factors that maintain epithelial stem cell function. This loss was not restored during MH, suggesting a sustained impairment of epithelial niche support. The top uniquely expressed gene defining this cluster was OGN. OGN has not previously been implicated in colon inflammation; however, it has been shown to functionally promote epithelial repair in COPD models of lung injury, raising the possibility that it serves a similar function in the colon and that its persistent loss may limit epithelial regeneration during MH ^61^.

Our spatial and protein-level analyses reveal a coordinated shift between fibroblast states, characterized by loss of OGN-expressing SCN fibroblasts and expansion of PDPN⁺ inflammatory fibroblasts. This transition reflects a broader reprogramming of the stromal compartment, in which homeostatic, niche-supporting functions are replaced by inflammatory and matrix-remodeling programs. Such an imbalance is likely to impair epithelial regeneration and limit restoration of normal tissue architecture during MH, consistent with the persistent crypt abnormalities observed by SHG imaging.

Beyond the fibroblast compartment, our spatial and transcriptomic analyses reveal marked plasticity within the pericyte population. While fibroblasts are well established as key mediators of inflammation and wound healing in the intestinal mucosa, the role of pericytes outside their traditional vascular context remains less well defined. Under homeostatic conditions, pericytes function as mural cells tightly associated with the endothelium, contributing to vascular stability. During active UC, however, we observed spatial decoupling of pericytes from the vasculature, accompanied by a transition toward a matrix-remodeling phenotype. This process is consistent with a mural-to-mesenchymal transition, in which pericytes detach from the vascular niche and redistribute into the lamina propria ^62-65^. Such redistribution may contribute to vascular dysfunction and increased permeability characteristic of IBD ^66,67^. Similar pericyte reprogramming has been described in cancer, where we and others have identified matrix-remodeling pericyte populations (matriPer^41^) across multiple tumor types, suggesting that this transition reflects a conserved stromal response across pathological contexts ^62,68,69^. This reprogrammed pericyte state is only partially resolved during MH, indicating a sustained contribution to ECM remodeling long after the inflammatory trigger has subsided.

At the tissue scale, our spatial transcriptomic analysis revealed reorganization of the colonic mucosa at the level of anatomical domains. We observed pronounced structural changes across disease states, accompanied by domain-specific transcriptional reprogramming. For example, the lamina propria exhibited a shift from transcriptional programs supporting crypt-niche maintenance in healthy tissue, to immune activation during active disease, and to ECM and collagen assembly during MH.

The GALT also underwent profound structural disassembly during active colitis, characterized by spatial redistribution of stromal and immune populations. During MH, however, the GALT did not fully return to homeostasis and instead remained compositionally altered. These changes are consistent with ongoing antigenic stimulation and localized immune activation. More broadly, these observations indicate that domain-level organization of the mucosa is not fully restored following inflammation, supporting the concept that MH represents a structurally reassembled but functionally reprogrammed tissue state.

The clinical management of UC is fundamentally limited by high rates of therapeutic failure. Despite an expanding arsenal of targeted biological therapies, up to 30-40% of patients exhibit primary non-response to biologics and small molecules, while a substantial fraction of initial responders eventually lose response over time ^21^. Identifying predictive biomarkers to guide therapeutic interventions and avoid prolonged exposure to ineffective treatments therefore remains a critical unmet clinical need. In this context, we demonstrate that elevated baseline signatures of inflammatory fibroblasts and pericytes robustly predict patient non-response to infliximab. These findings highlight a key limitation in current IBD management: while modern biologics effectively target specific immune pathways, they may not fully reverse the reprogramming of a heavily damaged stromal scaffold. Our findings also have implications for current therapeutic endpoints. Although endoscopic remission is the accepted treatment goal in UC, increasing evidence suggests that histologic healing provides a more stringent measure of tissue recovery and is associated with improved long-term outcomes^15^. Notably, most patients classified as being in MH in our cohort, as well as patients classified as being in remission in the external cohorts, fulfilled criteria for histologic healing, yet persistent stromal remodeling remained readily detectable. These observations suggest that resolution of endoscopic and histologic inflammation may not be sufficient to restore tissue homeostasis. Instead, our data raise the possibility that stromal healing represents an additional dimension of tissue recovery, characterized by restoration of stromal cell states, tissue architecture, and epithelial**-**stromal interactions.

Collectively, our work demonstrates that despite the macroscopic appearance of a healthy mucosa, patients with UC continue to harbor a persistent stromal "inflammatory memory." This state is characterized by sustained compositional shifts across multiple intestinal compartments and enduring alterations in stromal composition and architecture. In this context, stromal healing may represent an additional dimension of tissue recovery beyond endoscopic and histologic healing. Therapeutic strategies aimed at resolving pathogenic inflammatory fibroblast states, regenerating the OGN⁺ stem cell niche, and re-establishing pericyte-endothelial coupling may enable more complete mucosal recovery and reduce the risk of relapse in UC.

## Materials and method

### Ethics statement

All clinical data were collected following approval by Rabin and Schneider Children’s Medical Centers (IRB protocols #RMC-0992-20, RMC-0411-21, RMC-0799-21) and the Weizmann Institute of Science (IRB #2325-2) Institutional Review Boards.

### Human Patient Samples

Mucosal biopsies were harvested via colonoscopy from the rectosigmoid colon to ensure anatomical consistency and minimize spatial heterogeneity. These samples were obtained from patients treated at Schneider Children’s Medical Center of Israel and Rabin Medical Center (Beilinson Hospital; informed consent to study the tissue was obtained via Schneider and Rabin Medical Centers (IRB protocols #RMC-0992-20, #RMC-0411-21, #RMC-0799-21), and the Weizmann Institute of Science (IRB protocol #2325-2) Institutional Review Boards. The cohort included a total of 89 individuals, 59 recruited at Schneider Children’s Medical Center and 30 at Rabin Medical Center. This cohort comprised 28 healthy controls, 37 patients with active UC, and 24 patients in endoscopic remission (MH). The vast majority (75%) of the MH group had concomitant histological remission (Nancy score= 0; see Supplementary Table 1 for details). Multiple biopsies were harvested from the same region and processed immediately. Samples were either fixed in PFA, frozen at -80°C, or preserved in Tissue Storage Solution (Miltenyi Biotec, Cat# 140-005-795).

### Cell line

The human colon-derived fibroblast cell line CCD18-Co was purchased from ATCC. CCD18-Co cells were expanded and cryopreserved according to ATCC guidelines. Cells were cultured in Eagle’s Minimum Essential Medium (EMEM; Sigma Aldrich) supplemented with 10% Fetal Bovine Serum (FBS; Glibco^TM^, ThermoFisher Scientific), 1% Penicillin and Streptomycin (P/S; BI-Biological Industries), and 1% L-Glutamine (BI-Biological Industries). Cells were routinely monitored for mycoplasma infections.

### Human colon tissue dissociation for single cell RNA sequencing

Biopsy samples were rinsed in cold cell-storage-solution (Miltenyi Biotec, 140-005-795) to remove debris. To eliminate mucus and epithelial cells, tissues were incubated in 20 mL Hank’s Balanced Salt Solution (HBSS) without calcium and magnesium, with 5 mM Ethylenediaminetetraacetic acid (EDTA) and 1% P/S and placed in a shaking air bath at 37°C (250 rpm) at a 45° angle for 15 min. Supernatant was discarded, and the incubation was repeated with fresh EDTA solution for another 15 min. Following chelation, tissues were washed in 20 mL ice-cold Phosphate Buffered Saline (PBS) without calcium and magnesium. For enzymatic digestion, tissues were transferred to gentleMACS™ C Tubes (Miltenyi Biotec) containing 3 mL digestion medium (RPMI 1640 with 10% FCS, 1% L-glutamine, 1% HEPES, 1% Sodium Pyruvate, 1% P/S, 20 mg/mL Collagenase D, 2.5 mg/mL Liberase, and 10 mg/mL DNase I). Mechanical and enzymatic dissociation was carried out using the gentleMACS™ Dissociator (Miltenyi Biotec) and quenched by 15 mL of MACS buffer (PBS with 0.5% Bovine Serum Albumin (BSA) and 2 mM EDTA). The suspension was passed through a 70-µm cell strainer and centrifuged at 350 × *g* for 5 min at 4°C. To remove erythrocytes, the pellet was resuspended in 1 mL of red blood cell lysis solution and incubated on ice for 5 minutes. The reaction was Neutralized with 8 mL MACS buffer, followed by centrifugation at 350 × *g* for 5 min at 4°C. The final cell pellet was resuspended in fixation buffer in accordance with the Chromium Single Cell Gene Expression Flex protocol (10x Genomics, CG000527 Rev F). Cells from each patient were fixed with formaldehyde at 4°C overnight, following the manufacturer’s instructions. After buffer exchange, the fixed cells were stored at –80°C until subsequent processing for downstream library preparation.

### Single-Cell Fixed RNA Profiling of Multiplexed Samples for Gene Expression

The scRNA-seq dataset was generated from 15 fixed patient biopsies, comprising 5 healthy, 6 active UC, and 4 MH samples. An additional MH sample (ECM_034) was sequenced but excluded from downstream analysis due to poor library quality. Samples were processed according to the chromium fixed RNA profiling for multiplexed samples protocol (10x Genomics, CG000527 Rev F). Briefly, fixed cells were hybridized with the human transcriptome probe set for 24 h at 42°C. Post-hybridization, samples were pooled and loaded onto the Chromium X Controller for droplet generation across four independent capture reactions. Library preparation, including probe ligation and amplification, followed manufacturer’s instructions. Libraries were loaded onto Illumina NovaSeq X platform using a 1.5B flow cell at a final loading concentration of 180 pM with a 10% PhiX spike-in. Raw data were processed using Cell Ranger v8.0.0 (10x Genomics).

### Single cell analysis

Single-cell RNA-seq data from 15 human samples was analyzed using the Seurat (v5) pipeline^70^. Raw count matrices generated from 10x Genomics libraries were imported using Read10X, and individual samples were merged into a unified Seurat object. Quality control filtering retained cells with 200–10,000 detected genes and mitochondrial transcript content below 20%, thereby excluding low-quality cells and mitigating potential doublets. Gene expression values were log-normalized (scale factor 10,000), and 2,000 highly variable genes were identified using the VST method, followed by data scaling and principal component analysis (PCA) using the top 20 components. To correct for inter-patient variability, Harmony was applied using patient identity as the integration variable, and the corrected embeddings were used for nearest-neighbor graph construction, clustering, and visualization. Cells were clustered using a shared nearest neighbor (SNN) graph and the Louvain algorithm. Dimensionality reduction was performed using UMAP based on the Harmony embeddings. Patient metadata, including disease state (healthy, active UC and MH), were integrated by matching sample identifiers. Cell type composition across disease states was quantified by calculating relative cell proportions per sample and aggregating these values by disease group. Differences in cell type composition between disease groups were assessed using the Kruskal-Wallis test.

### U method analysis

Cell type marker genes were identified using the U-method - a probability-based framework for identifying uniquely expressed genes (UEGs) within any clustering of scRNA-seq data. For each gene and each cluster, the method compares the detection probability in that cluster with the highest detection probability observed in any other cluster (Umethod R package). A U-score threshold ≥ 0.2 and minimum within-cluster expression frequency (*P*_*in*_ ≥ 0.5) were applied to define UEGs. Clusters exhibiting features consistent with doublet artifacts, cellular stress, actively dividing or exceptionally small sizes were identified during the analysis. As these clusters lacked distinct functional signatures, they were excluded from the U-method *P-out* calculation to avoid skewing marker identification, but were retained in global visualizations and dot plots to provide a comprehensive overview of the dataset.

### Cell-Cell Communication Analysis

Ligand-receptor mediated signaling interactions were inferred from scRNA-seq data using CellChat (v2.1.2) ^37^. Analysis was performed on 15 patients across three clinical groups (Healthy, Active UC, and MH), with CellChat applied independently to each patient to enable patient-level resolution. For each sample, raw counts were extracted from the Seurat object and a CellChat object was constructed using normalized expression data, with cell type annotations derived from unsupervised clustering, comprising eleven cell populations: Epithelial, T cells, Plasma, B cells, Fibroblasts, Monocytes, Endothelial, Granulocytes, Pericytes, Glia, and Enteroendocrine cells. The CellChatDB.human database was used as the reference for ligand-receptor interactions. Overexpressed genes and ligand-receptor pairs were identified using default parameters, followed by computation of communication probabilities and pathway-level communication scores. Cell populations with fewer than 10 cells per interaction or fewer than 50 cells per patient were excluded from downstream analysis. For each patient, aggregated interaction networks were computed, and sender and receiver interaction strengths were derived from the aggregate weight matrix. Group-level comparisons were performed by averaging per-patient sender and receiver scores within each clinical group. Statistical differences across groups were assessed using the Kruskal-Wallis test.

### Crypt Zonation of Stromal subsets from Single-Cell RNA-seq analysis

Upper and lower crypt regions were analyzed using the AUCell R package^71^ to score individual cells for gene signatures. The crypt-bottom signature comprised essential Wnt/R-spondin mediators (*GREM1, RSPO3, IGF1, WNT2, WNT2B, VEGFD*), while the crypt-top signature was defined by BMP-associated genes (*PDGFRA, SOX6, BMP2, BMP3, BMP4, BMP5, BMP7, WNT5A, NRG1*)^25^.

### Second harmonic generation

#### Second harmonic generation imaging

Distal FFPE colon samples were imaged such that the entire section in a horizontal orientation (excluding artifact- and fold-containing areas) was captured in separate fields of view (FOVs) for subsequent analysis, using SHG microscopy on an upright Leica TCS SP8 MP microscope equipped with external non-descanned detectors (NDD) HyD and an Acusto Optical Tunable Filter (Leica microsystems CMS GmbH, Germany). SHG signals were excited by an 885nm laser line from a tunable femtosecond laser - Coherent vision II, 680-1080 nm; (Coherent GmbH USA). Emission was collected *via* an external NDD HyD detector using a 440 nm long pass filter. The transmitted signal was collected using a PMT detector in transmission position to capture overall morphology. DRAQ5 staining was imaged using a HeNe 633 laser with emission collected between 670-760nm. Images were acquired at 2048 x 2048 resolution using an HC FLUOTAR L 25X/0.95 W VIS objective and with the following settings: Scan Speed 600 Hz; Zoom 0.75; Line averaging 3 and Bit depth 16. The field of view was 590.48 x 590.48 µm with a pixel size of 0.29 µm. Z stacks were acquired using a galvo stage with a step size of 5.11 µm and 0.57 µm intervals. Images were processed and visualized using LASX software (Leica Application Suite XLeica microsystems CMS GmbH).

#### Second harmonic generation analysis

We started with Max projection of the 3D stack, and trained Ilastik (1.3.3b2)^72,73^ AutoContext Pixel Classifier to classify pixels into crypts vs. collagen fibers, on selected images. A dedicated Fiji macro applied this classifier to each image and segmented the individual crypts using Hysteresis Thresholding from 3D ImageJ Suite plugin^72^. Holes in the crypt objects were filled and objects smaller than 300 µm^2^ were discarded. For each image we measured the number of crypts completely included in the image, and the average size of those crypts. Images of segmented crypts color-coded by these measurements were created using MorphoLibJ plugin^74^. This automatic process segmented correctly most of the crypts. Further manual correction was done to fix for segmentation errors. All the measurements were recalculated based on the corrected segments. The macro allows for this by saving crypts contours in a file, and by providing an update mode to calculate the measurements from a segments file. The macro is available on Github DOI 10.5281/zenodo.4172576 (https://github.com/WIS-MICCCellObservatory/Crypts_SpatialOrganization/).

### Visium HD spatial transcriptomics

The Visium HD dataset was assembled from 12 patients distributed equally across three groups: healthy controls, active UC, and MH. four colonoscopy biopsies were placed within a single capture area. Sections (4 µm thick) from FFPE tissue blocks were mounted onto Poly-Prep slides (Sigma-Aldrich, P0425-72EA) adhering to the protocols outlined in the Visium HD Tissue Preparation Handbook (CG000684 Rev A). H&E staining and imaging were performed according to the handbook. Brightfield whole-slide images were acquired using PhenoImager Fusion 2.0 upright scanner. Samples were then processed and libraries were constructed using the Visium HD Spatial Gene Expression Reagents Kit (CG000685 Rev B). Libraries were loaded onto individual lanes of an Illumina NovaSeq X platform using 150 pM in a 1.5B flow cell. Data processing was conducted using Space Ranger (version 4.0). All downstream analyses in this study were performed using the 8-µm binned data.

### Spatial Mapping of Single-Cell Derived UEG Signatures via Visium HD

Single cell derived gene signatures were projected onto Visium HD spatial transcriptomic data to map stromal cell populations *in situ*. Spots expressing signature-defined marker genes were identified and visualized, with each spot representing a spatial location positive for the given signature.

For GALT fibroblasts, an additional gating strategy was applied to increase specificity in this densely populated compartment. Spots were classified as GALT fibroblast-positive only if they co-expressed *CCL19* together with either MFAP4 or LUM, two fibroblast-specific UEGs identified in the single-cell analysis. This approach enriched for fibroblast-specific signal and minimized contribution from other non-fibroblast *CCL19*-expressing cells.

### Relative Risk (RR) analysis

Spatial enrichment analysis was performed to quantify the preferential localization of cell-type specific transcriptional programs within defined anatomical regions. Marker gene sets for each cell type were derived using the U-method (Uscore ≥ 0.25, minimum expression frequency ≥ 0.4, top 3 U-markers). For each Visium HD sample, spatial coordinates were extracted and normalized to a unit scale to ensure comparability across samples. A data-driven neighborhood radius was defined as a multiple (k-steps, k=1) of the median nearest-neighbor distance, enabling adaptive scaling to tissue-specific spot density. For each region of interest (e.g., GALT), spots annotated by Loupe-derived clustering were used as reference points, and all neighboring spots within the normalized radius were classified as proximal. Cell-type specific enrichment was then quantified using a “pure marker” framework, in which spots were assigned to a given cell type if they expressed at least one marker from that set and none from other cell-type marker sets. For each cell type, enrichment was computed as a relative risk (RR), defined as the ratio between the probability of observing pure marker-positive spots within the defined spatial radius and the corresponding probability outside the radius. Log-transformed RR values were used for comparative analyses across disease states.

### Human Bulk RNA-sequencing

For transcriptomic analysis, fresh-frozen colonic biopsy specimens (average size 2x2 mm) from 43 patients (12 healthy controls, 20 active UC, and 11 MH) were utilized. Tissues were thawed in 400 μL Tri-reagent (Sigma-Aldrich) and mechanically homogenized via bead beating, followed by a short centrifugation step to pull down the beads and tissue debris. Library preparation was performed using the mcSCRB-seq protocol ^75^ with minor modifications. Reverse transcription was applied on up to 500 ng of RNA, with a final volume of 20 μL (1×Maxima hour Buffer, 1 mM dNTPs, 2 μM TSO* E5V6NEXT, 7.5% PEG8000, 20U Maxima H enzyme, 2 μL barcoded RT primer). Subsequent steps were applied as mentioned in the protocol. Library preparation was performed using Nextera XT kit (Illumina) on 1-2.5 ng amplified cDNA. Library final concentration was 1.8-2 nM, and sequencing was done using the Novaseq 6000/X (Illumina) sequencing machine aiming at 40 M reads per sample with the following settings: Read1- 16bp, Index1-10bp, Index2-10bp, Read2-66bp.

### Human Bulk RNA-sequencing analysis

Illumina output files were demultiplexed and aligned to the human reference genome (GRCh38) using UTAP with CUTADAPT default trimming parameters. Mitochondrial, ribosomal, and non-protein-coding genes were excluded from downstream analysis. Protein-coding genes were defined using Ensembl BioMart annotation (GRCm38, release 91). Gene expression was normalized per sample by the sum of UMIs of genes individually contributing less than 5% of the total sample UMI count. Samples expressing fewer than 1,500 genes after filtering were excluded. Gene signatures for the stromal subsets were defined using the top 10 UEGs. The SCN fibroblast signature was derived from UEGs identified within the stromal subsets analysis. For inflammatory fibroblasts and pericytes, UEGs were identified by comparing these subpopulations to all other cell lineages in the full scRNA-seq dataset (Supplementary Table 8), to maximize specificity and minimize inclusion of genes expressed in non-stromal populations.

### Human Bulk RNA-seq Analysis validation cohorts

Stromal signatures were evaluated across five independent publicly available bulk transcriptomic and microarray datasets retrieved from the NCBI Gene Expression Omnibus (GEO). The inflammatory fibroblast, pericyte, and SCN fibroblast signatures were applied consistently across all cohorts. Across these datasets, patients classified as treatment responders or in MH were generally defined by a combination of endoscopic mucosal healing and histological remission, with the latter predominantly determined by a Geboes^76^ or histological score of ≤ 1. In the GSE128682 cohort (RNA-seq) samples were stratified into healthy controls, active UC, and MH. For consistency across our analyses, the endoscopically defined "remission" group from the original metadata is referred to herein as MH. In the GSE16879 cohort (Affymetrix microarray), UC and CD-colon patients treated with infliximab were stratified into responder and non-responder groups based on treatment response metadata; analyses were performed at baseline and post-treatment timepoints. In the GSE73661 cohort (microarray), UC patients treated with vedolizumab (VDZ) or infliximab (IFX, assessed at Week 4/6) were classified as responders or non-responders based on endoscopic sub-score and clinical annotation respectively; for VDZ-treated patients, Week 52 was selected as the endpoint since this is the time at which responding patients achieve MH. SCN fibroblast signatures were not evaluated in this cohort due to insufficient gene representation on the platform. Two additional microarray cohorts (GSE12251 and GSE14580) comprising pre-treatment biopsies and response data (but not post-treatment biopsies) from UC patients, were analyzed to assess baseline predictors of treatment response. For microarray datasets, in cases where multiple probes mapped to a single gene, their expression values were averaged to yield a single representative value per gene. In the GSE12251 cohort, as described in the original study, two pre-treatment biopsies were obtained from one patient, therefore, sample GSM307892 was removed to ensure unique patient representation. For all datasets, per-sample composite signature scores were derived by calculating the mean expression of the signature genes, followed by Z-score standardization across the dataset. Group differences were assessed using Wilcoxon rank-sum. To evaluate the predictive performance of the baseline signatures for treatment response, receiver operating characteristic (ROC) curves were generated. The area under the curve (AUC) was calculated for individual cell-type signatures, as well as for a combined predictive model constructed using logistic regression.

### In vitro inflammation assay

CCD-18Co cells were seeded into 6-well plates at a density of 0.3 x 10^6 cells per well in complete DMEM and allowed to adhere overnight. The following day, the culture medium was replaced with either fresh complete DMEM (control) or DMEM supplemented with 1 µg/mL TNF-α and 1 µg/mL IL-17 (human recombinant proteins, Peprotech), and cells were incubated for 24 hours at 37°C. Following incubation, the cells were lysed in TRIzol Reagent (Thermo Fisher), total RNA was extracted and assessed for quality and RNA-seq libraries were generated using a bulk adaptation of the MARS-seq protocol as previously described ^77,78^. The libraries were sequenced on a NovaSeq X Plus platform (Illumina, paired-end, 100-cycle kit) with a minimum of 10 million reads per sample. Raw reads were demultiplexed and aligned to the human reference genome (hg38) using UTAP with CUTADAPT default trimming parameters. One sample (sample 41) was excluded from downstream analysis due to insufficient read counts. Read counts were normalized and differential expression analysis was performed using the DESeq2 pipeline. For visualization, regularized log-transformed (rlog) expression values were Z-score normalized per gene.

### Formalin-fixed, paraffin-embedded (FFPE) blocks preparation

Patient samples designated for formalin-fixed paraffin-embedded (FFPE) processing were placed in 4% paraformaldehyde (PFA) immediately following colonoscopic biopsy, subsequently transferred to 1% PFA, and embedded in paraffin blocks.

### Multiplexed immunofluorescent (MxIF) staining

5-μm-thick sections were deparaffinized in xylene and incubated in 10% neutral buffered formalin (prepared by 1:25 dilution of 37% formaldehyde solution in PBS). Antigen retrieval was performed using citrate buffer (pH 6.0) for the epithelial cell cocktail (CK18, CK7, EpCAM), PDGFRβ, MCAM, and CD31; or Tris-EDTA buffer (pH 9.0) for PDPN, RGS5 and CD45. Slides were then blocked using 10% BSA in PBS (BlockerTM BSA, Thermo Scientific). All primary antibodies (Supplementary Table 12) were diluted in 2% BSA PBS with 0.05% Tween (PBST) and used in a multiplexed manner using the OPAL reagents (Akoya Biosciences). Primary antibody incubation overnight at 4C was followed by incubation with the complementary horseradish peroxidase (HRP) conjugated secondary antibody. Next, the slides were incubated with Opal reagents and subsequently washed to perform antigen retrieval as detailed above. Then, slides were incubated with the subsequent primary antibody or with DAPI at the end of the MxIF staining cycle. We used the following staining sequences: PDGFRβ → Epithelial cell cocktail → OGN → PDPN → DAPI for Figure 3K; CD31 → Epithelial cell cocktail → RGS5 → PDGFRβ → MCAM → CD45 → DAPI for Figure 4I. Each antibody was validated and optimized separately and then MxIF was optimized. The slides for Figure 3K were imaged with a PhenoImager HT scanner (Akoya Biosciences) and the slides for Figure 4I were imaged with a PhenoImager Fusion 2.0 upright scanner (Akoya Biosciences).

### MxIF Imaging analysis

Background removal was performed using the Akoya spectral unmixing algorithm with a non-stained reference slide. Images were analyzed using QuPath (version 0.3.4 and version 0.5.1 RRID:SCR_018257), and ImageJ-Fiji ^79^ for snapshots and single-channel representative images. To ensure consistent and reliable spatial analysis, regions of interest (ROIs) exhibiting a uniform transverse tissue orientation were selected from each sample. Object-based analysis was performed following nuclear segmentation using DAPI staining and the cellpose algorithm^80^ with the pretrained “nuclei” model. Cell classification was carried out using the “Random Trees” classifier trained to identify marker-positive cells based on marker-specific fluorescence intensity morphological features, DAPI signal, and autofluorescence channels. Classifiers were trained separately for each marker and subsequently combined into a composite classifier applied across all images. Marker positivity was quantified as the percentage of positive cells per tissue ROI relative to the total number of cells in that ROI.

Cellular subpopulations were classified based on specific marker combinations (Figures 3K and 4I). In both figures, epithelial cells were defined by positive staining for the epithelial marker cocktail (CK18, CK7, EpCAM). In Figure 3K, inflammatory fibroblasts were defined by double positive staining for PDGFRβ^+^ (general stromal marker in the colon) and PDPN^+^. As OGN is a secreted protein, its abundance was quantified using a pixel-based classifier in QuPath. In Figure 4I, endothelial cells were defined by CD31 staining and pericytes were identified based on MCAM and RGS5 expression. While PDGFRβ is also known to be a pericyte marker, it exhibited broad, non-mural specific staining within the mucosal stroma and was therefore excluded from the pericyte classification.

### Statistical analysis

Computational and statistical analyses were performed using R software (version 4.3.1, R Foundation for Statistical Computing, Vienna, Austria), Python (version 3.11.14), and Prism version 9.1.1 (GraphPad Software, USA). The specific statistical tests applied for each experiment are detailed in the respective Figure legends. Briefly, for normally distributed data, comparisons among more than two groups were performed using a one-way analysis of variance (ANOVA) followed by Tukey’s post-hoc correction. For data that did not follow a normal distribution, the non-parametric Wilcoxon rank-sum test was used for two-group comparisons, and the Kruskal-Wallis test was employed for comparing three or more groups. The association between two continuous variables was evaluated using Pearson’s correlation coefficient. Where applicable, correction for multiple hypothesis testing was performed using the Benjamini-Hochberg (BH) false discovery rate (FDR) procedure. Statistical significance was defined as a p-value or FDR < 0.05. In all Figures, significance levels are denoted as follows: *p < 0.05, **p < 0.01, ***p < 0.001.

## Data availability

All datasets generated during this study have been deposited in the Gene Expression Omnibus (GEO).

Publicly available datasets analyzed in this study are available in the GEO repository under the following accession numbers: GSE128682, GSE16879, GSE73661, GSE12251, and GSE14580.

Generative AI tools (OpenAI ChatGPT and Google Gemini) were used to assist with code debugging, code documentation, and implementation of small analysis functions based on pseudocode designed by the authors. All generated code was independently reviewed, tested, revised where necessary, and validated by the authors prior to use in the study, and the authors take full responsibility for the final content of the work.

## Acknowledgments

RSS is incumbent of The Robert and Yadelle Sklare Professorial Chair in Biochemistry. This study was supported by the Israel Science Foundation Personalized Medicine Program (IPMP) grant no. 3663/21, the Israel Cancer Research Fund (ICRF), and the Pfizer Emerging Science Fund.

## Lead contact

Further information and requests for resources and reagents should be directed to the Lead Contact, Ruth Scherz-Shouval.

## Declaration of generative AI and AI-assisted technologies in the manuscript preparation process

During the preparation of this work, the author(s) used generative AI tools (OpenAI ChatGPT and Google Gemini) for light language editing. The author(s) reviewed and edited the output as needed and take full responsibility for the content of the published article.

## Supplementary Figures

**Supplementary Figure 1.**
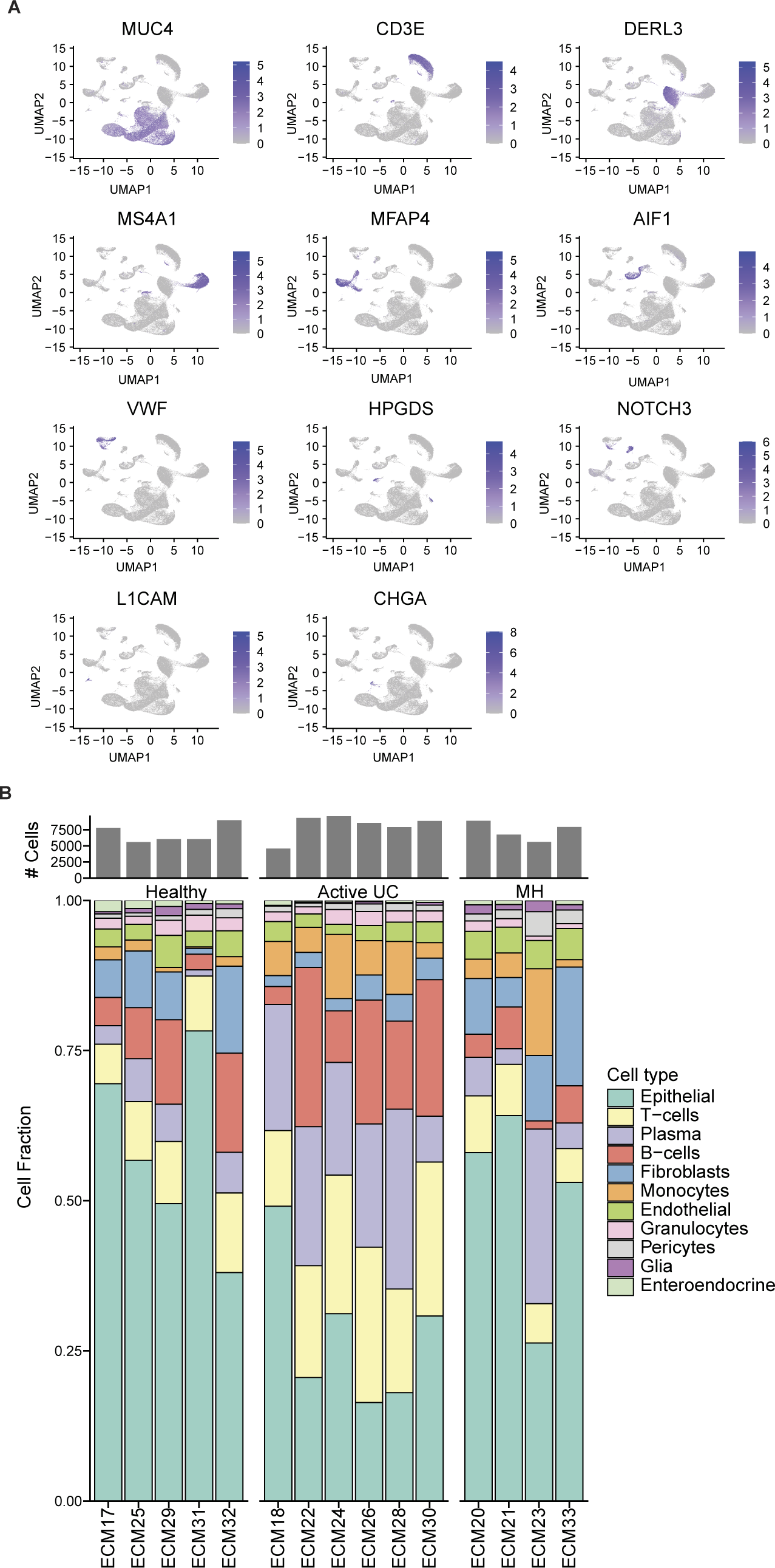
Cellular compositions of the scRNA-seq dataset. **(A)** Feature plots displaying log-normalized expression of the top UEGs defined using the U-method algorithm, and used to annotate the following cell lineages: Epithelial cells (*MUC4*), T-cells (*CD3E*), Plasma cells (*DERL3*), B-cells (*MS4A1*), Fibroblasts (*MFAP4*), Monocytes (*AIF1*), Endothelial cells (*VWF*), Granulocytes (*HPGDS*), Pericytes (*NOTCH3*), Glia (*L1CAM*) and Enteroendocrine cells (*CHGA*). **(B)** Patient-level cellular composition of the scRNA-seq dataset. The top bar plot displays the absolute number of cells analyzed per patient. The bottom stacked bar plot illustrates the fractional representation of each cell lineage per patient.

**Supplementary Figure 2.**
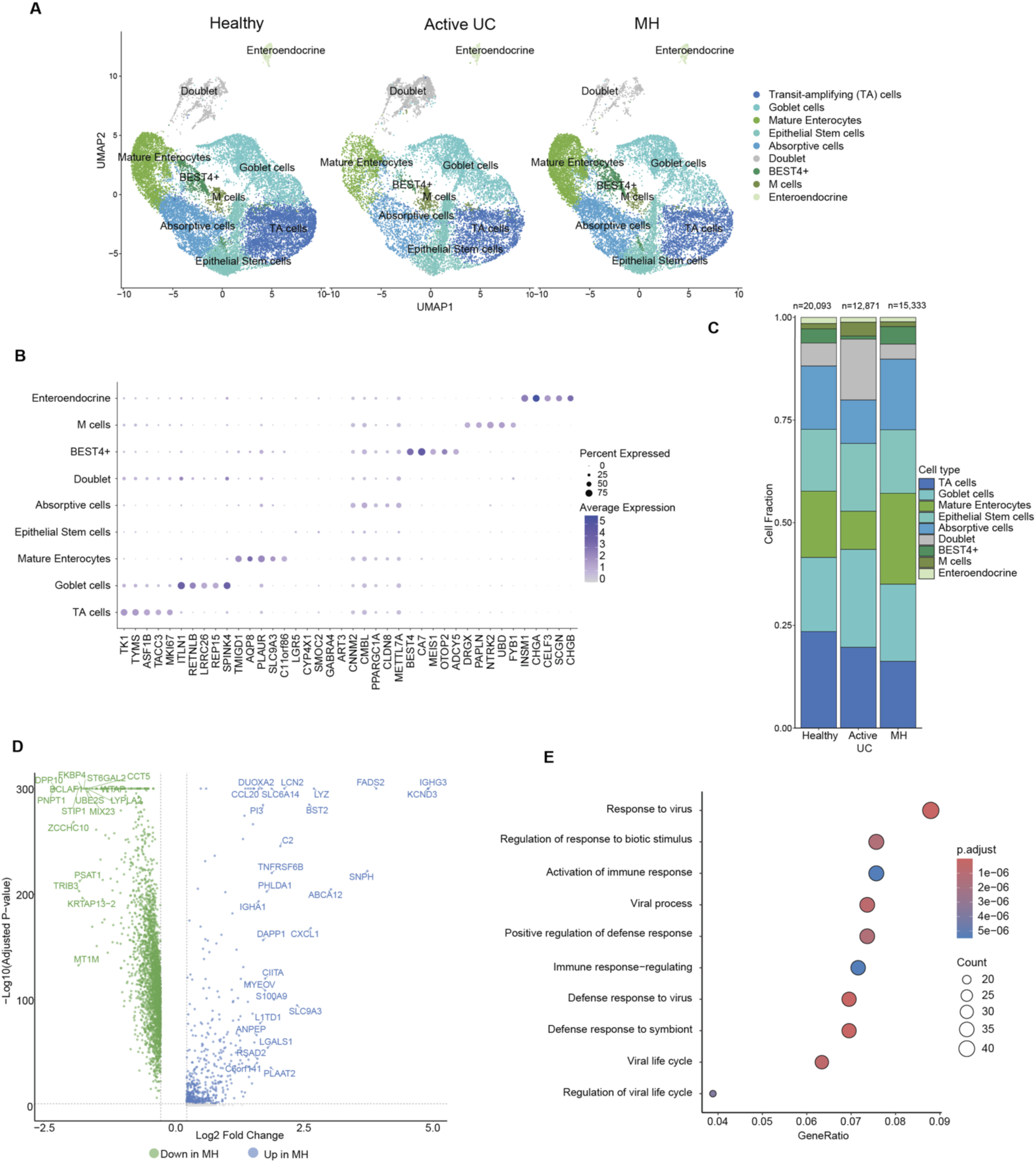
scRNA-seq of the colonic epithelial compartment reveals extensive compositional shifts across disease states. **(A)** Subclustering of epithelial cells using Seurat and UMAP visualization of the scRNA-seq dataset, split by disease state (Healthy, Active UC, and MH). **(B)** Dot plot showing the top 5 UEGs, defined using the U-method and used to annotate epithelial subpopulations. **(C)** Stacked bar plot illustrating the relative proportions of epithelial cell subsets across disease states. The total number of epithelial cells analyzed per condition is indicated**. (D)** Volcano plot displaying differentially expressed genes (DEGs) in the immune compartment of MH patients compared to healthy controls. Significantly upregulated (blue) and downregulated (green) genes in MH are highlighted. Highly significant genes are labeled (Fold Change ≥ 3 or ≤1/3, adjusted P≤ 0.005).. **(F)** Top 10 significantly enriched pathways for genes upregulated in the MH epithelial compartment, analyzed using clusterProfiler (adjusted Pvalue<0.05).

**Supplementary Figure 3.**
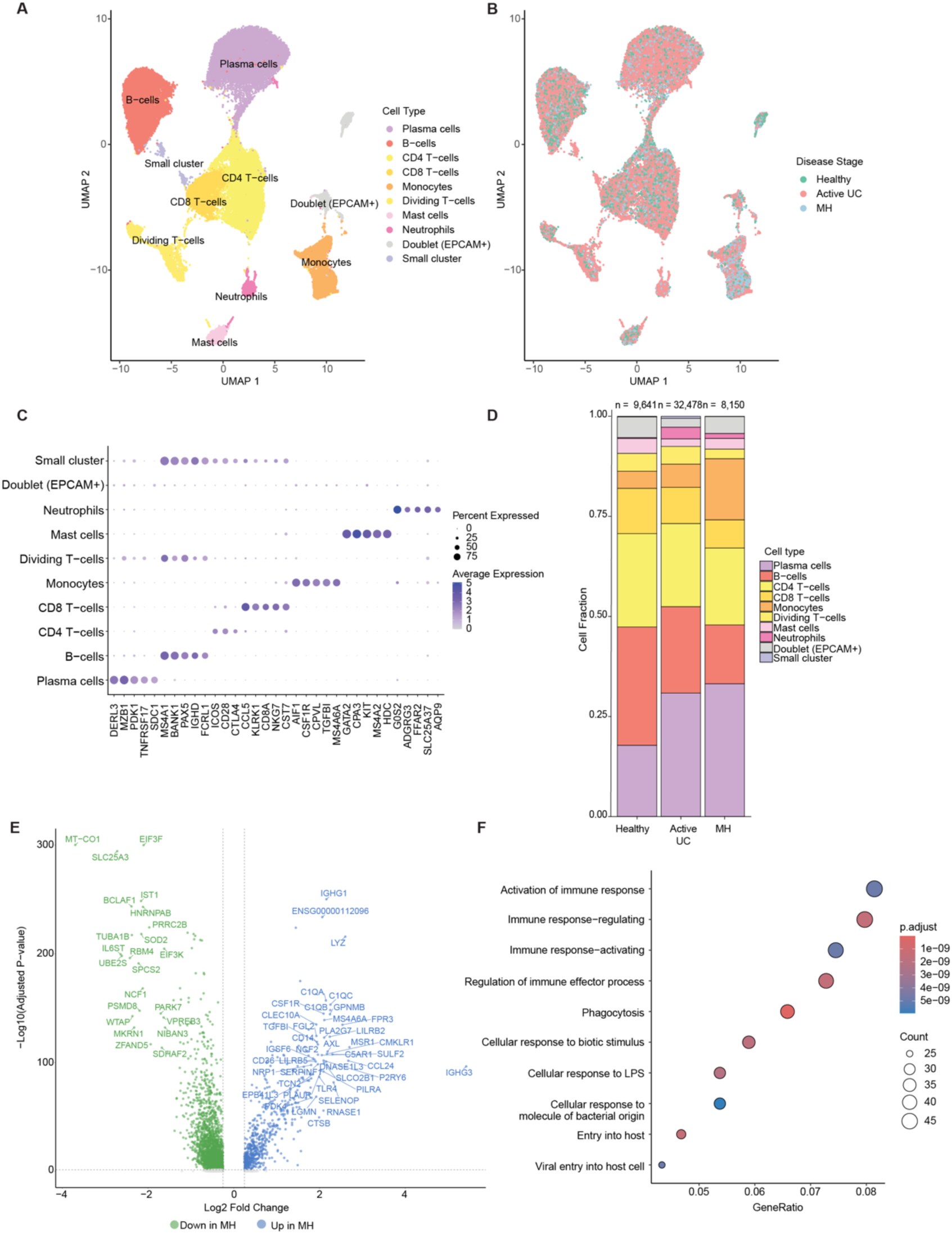
The immune compartment expands during active UC and exhibits persistent compositional alterations during mucosal healing. **(A-B)** Subclustering of the immune cell compartment using Seurat and UMAP visualization of the scRNA-seq dataset, colored by **(A)** major immune cell lineages and **(B)** clinical disease state (Healthy, Active UC, MH). **(C)** Dot plot showing the top 5 UEGs defined using the U-method and used to annotate immune subpopulations. **(D)** Stacked bar plot illustrating the relative proportions of immune cell subsets across disease states. The total number of immune cells analyzed per condition is indicated. **(E)** Volcano plot displaying differentially expressed genes (DEGs) in the immune compartment of MH patients compared to healthy controls. Significantly upregulated (blue) and downregulated (green) genes in MH are highlighted. Highly significant genes are labeled (Fold Change ≥ 3.5 or ≤1/3, adjusted P≤ 0.005). **(F)** Top 10 significantly enriched pathways for genes upregulated in the MH immune compartment, analyzed using clusterProfiler (adjusted Pvalue<0.05).

**Supplementary Figure 4.**
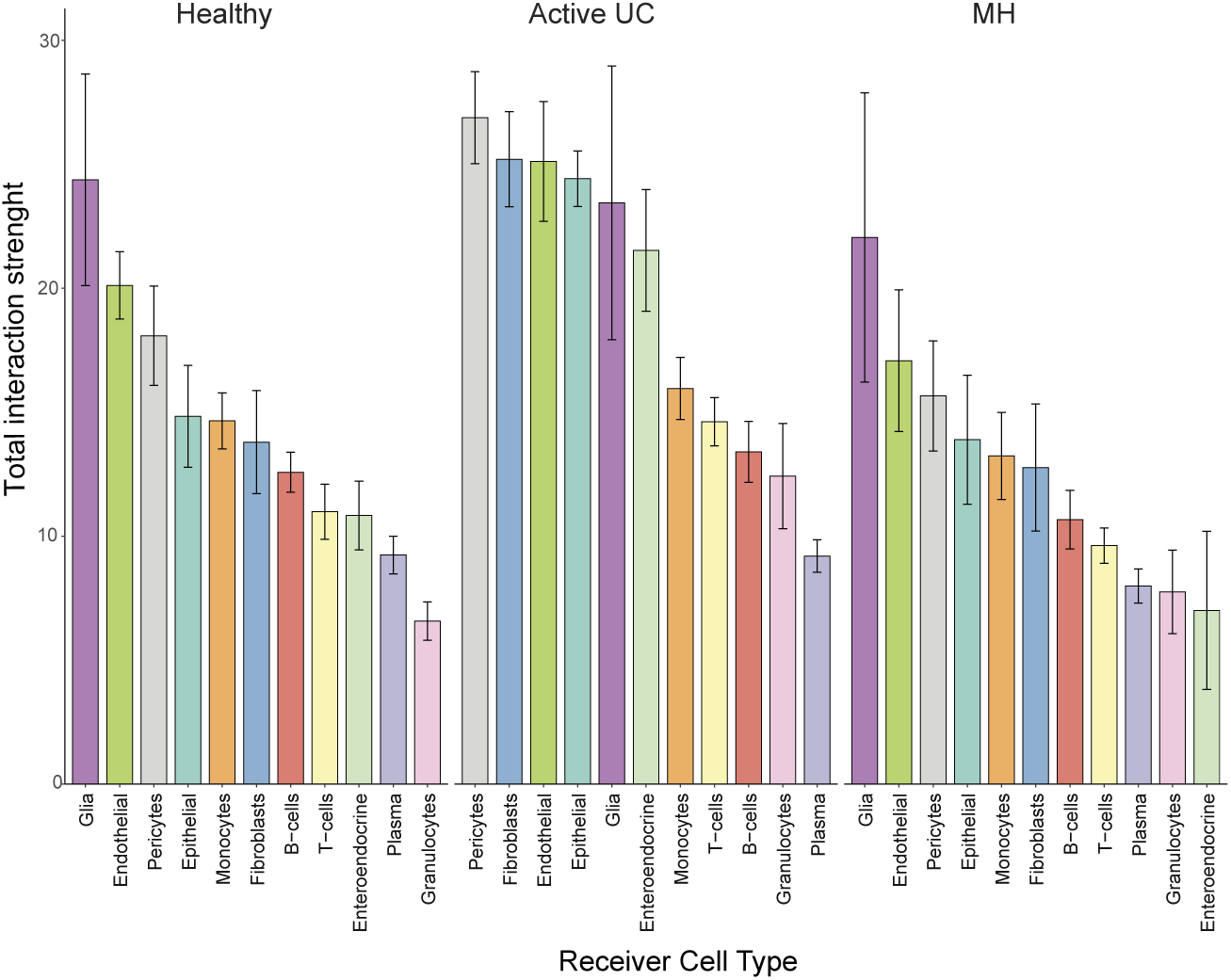
Cell-cell interaction patterns are altered across UC disease states. Bar plot representation of the signaling activity of each cell lineage based on cumulative ligand-receptor interaction scores, calculated using the CellChat algorithm applied to the scRNA-seq data described in Figure 1. Receiver signals are shown. Error bars represent mean ± SEM.

**Supplementary Figure 5.**
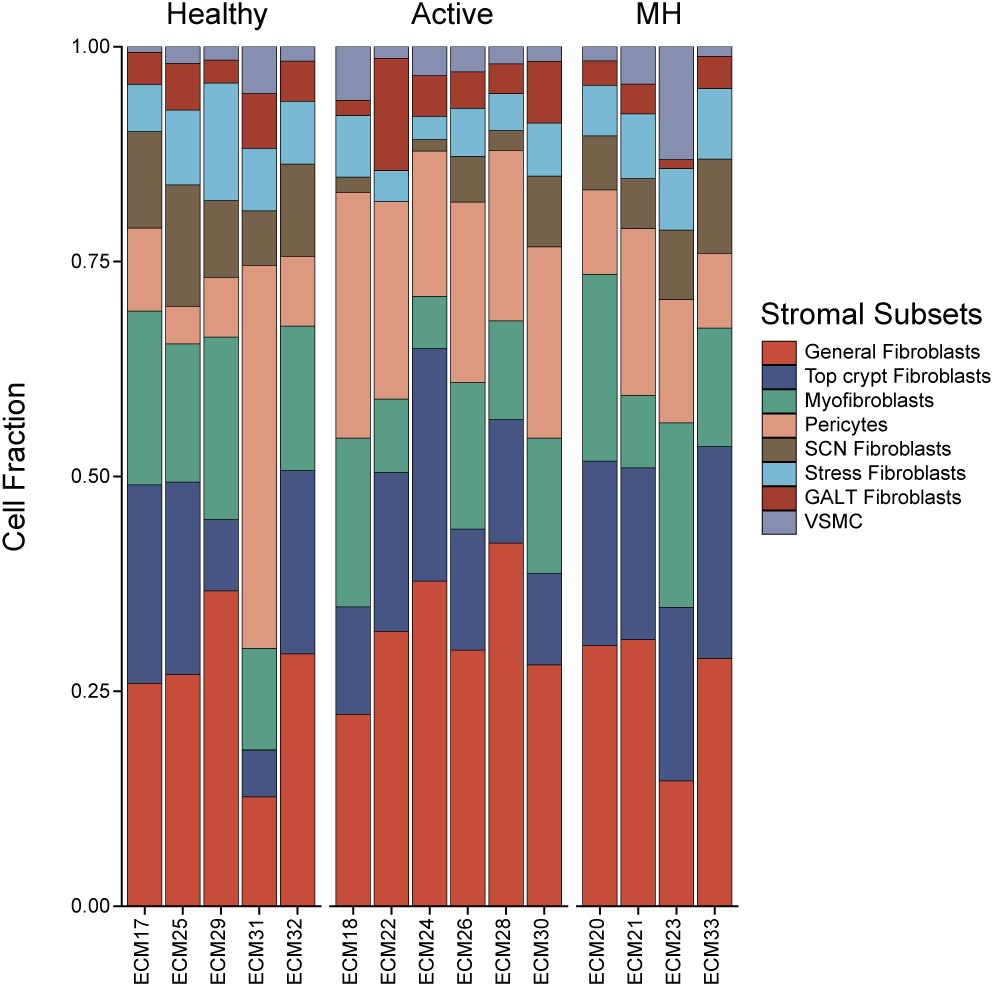
Stromal subpopulation dynamics across patients. Stacked bar plot illustrating the relative proportions of stromal subpopulations for individual patients across disease states, based on the scRNA-seq analysis shown in Figure 2A.

**Supplementary Figure 6.**
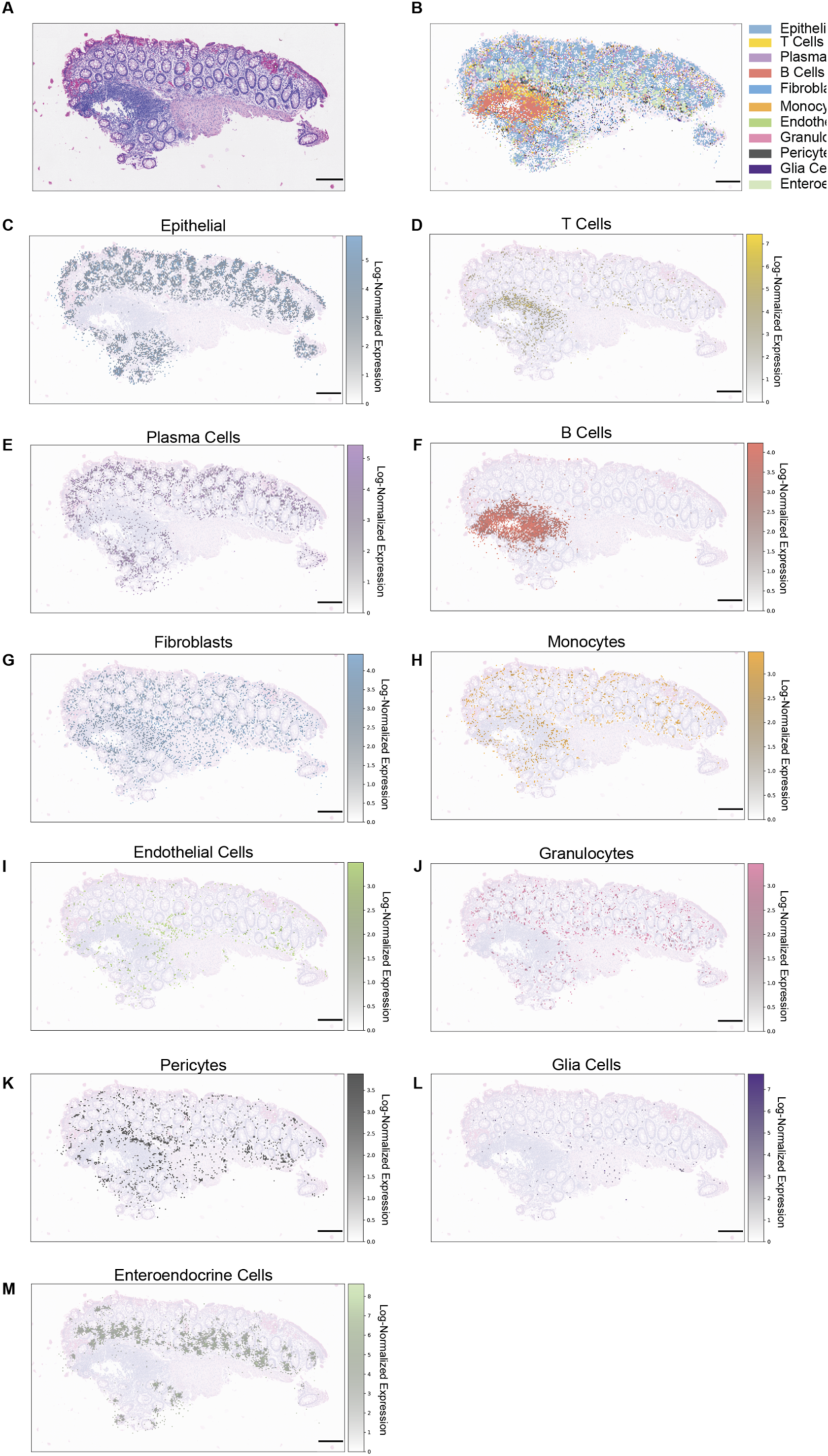
Spatial mapping of the major colonic cell lineages in a healthy patient using Visium HD. Visium HD spatial transcriptomics was performed on 12 biopsies as described in Figure 2E. **(A)** Representative H&E staining of the healthy colonic tissue section. **(B)** Composite spatial map illustrating the comprehensive distribution of all major cell lineages across the colonic architecture, based on the Visium HD analysis. Spot identity was determined by calculating a lineage-specific expression score, defined as the log-transformed sum of the top UEGs for each cell type. Each spot was then assigned to and colored by the cell lineage exhibiting the highest dominant expression score. **(C-M)** Individual Visium HD spatial feature plots showing the localized expression signatures of distinct colonic cell lineages. Spots are marked by the log-normalized expression of their top two UEGs: Epithelial (*MUC4*, *C19orf33*), T cells (*CD3E*, *CD3D*), Plasma cells (*DERL3*, *MZB1*), B cells (*MS4A1*, *PAX5*), Fibroblasts (*MFAP4*, *LUM*), Monocytes (*AIF1*, *CSF3R*), Endothelial cells (*VWF*, *CDH5*), Granulocytes (*HPGDS*, *CPA3*), Pericytes (*NOTCH3*, *RGS5*), Glia (*L1CAM*, *PLP1*), and Enteroendocrine cells (*CHGA*, *INSM1*).

**Supplementary Figure 7.**
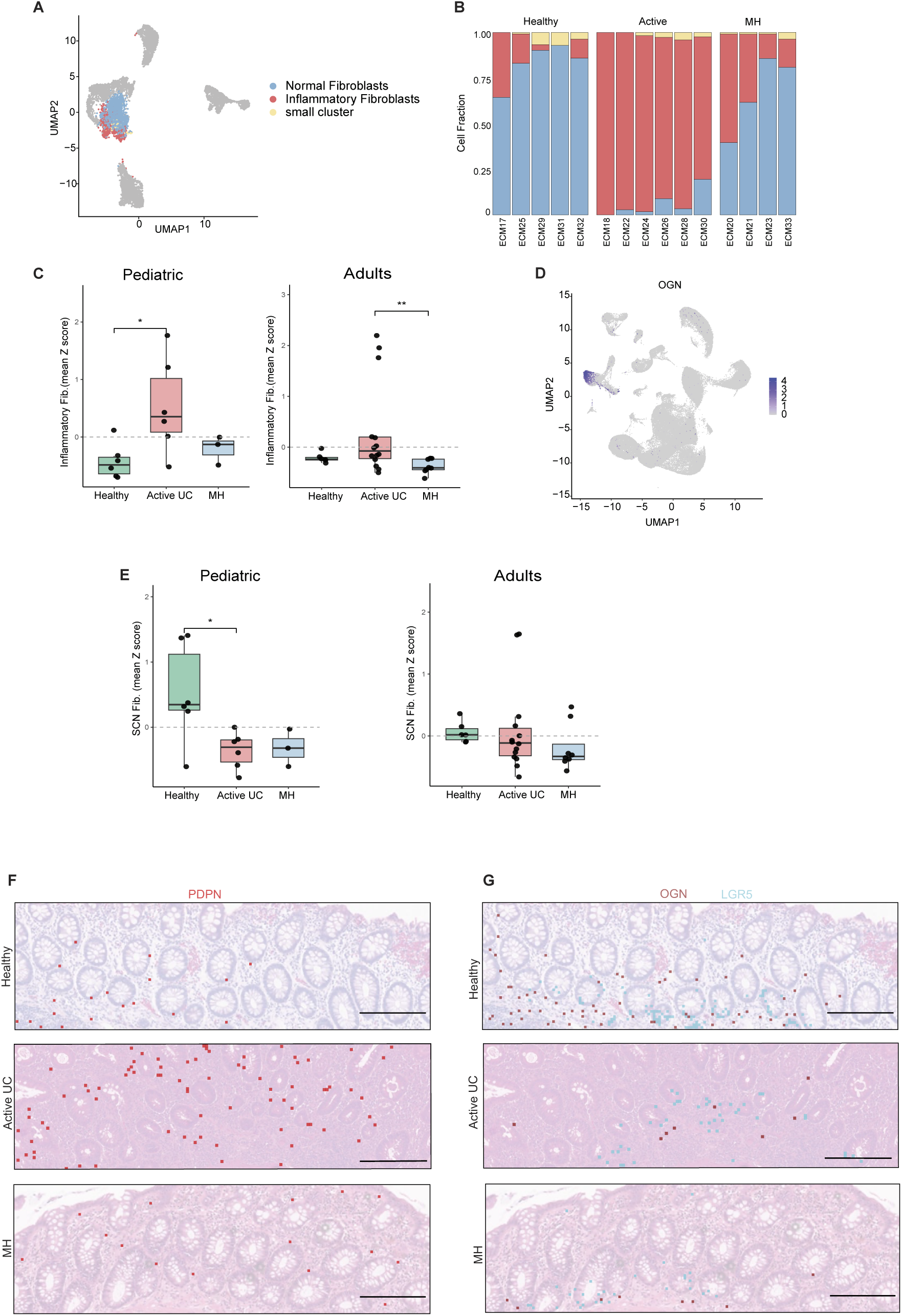
Transcriptomic and spatial profiling of inflammatory and SCN fibroblasts reveals dynamic changes across disease stages. **(A)** UMAP visualization of high-resolution General Fibroblast subsets: Normal (blue), Inflammatory (red), and minor (yellow), projected onto the broader stromal map (grey). **(B)** Stacked bar plot showing the relative proportions of General Fibroblast subpopulations across disease states and individual patients. **(C)** Mean Z-scores of the inflammatory fibroblast signature evaluated in pediatric (left) and adult (right) patients across disease states. Statistical significance is indicated (* p < 0.05; Wilcoxon test). **(D)** Feature plot displaying log-normalized expression of *OGN*, the top UEG defining the SCN subset, across all cell populations in the scRNA-seq dataset presented in Figure 1B. **(E)** Mean Z-scores of the SCN fibroblast signature evaluated in pediatric (left) and adult (right) patients across disease states. Statistical significance is indicated (** p < 0.01; Wilcoxon test). **(F-G)** Representative Visium HD spatial transcriptomic maps from different UC states overlaid on H&E sections, showing all spots positive for the Inflammatory fibroblast marker *PDPN* **(F)** and the SCN fibroblast marker *OGN*, co-mapped with the epithelial stem cell marker *LGR5* **(G)**. Scale bar - 200 µm.

**Supplementary Figure 8.**
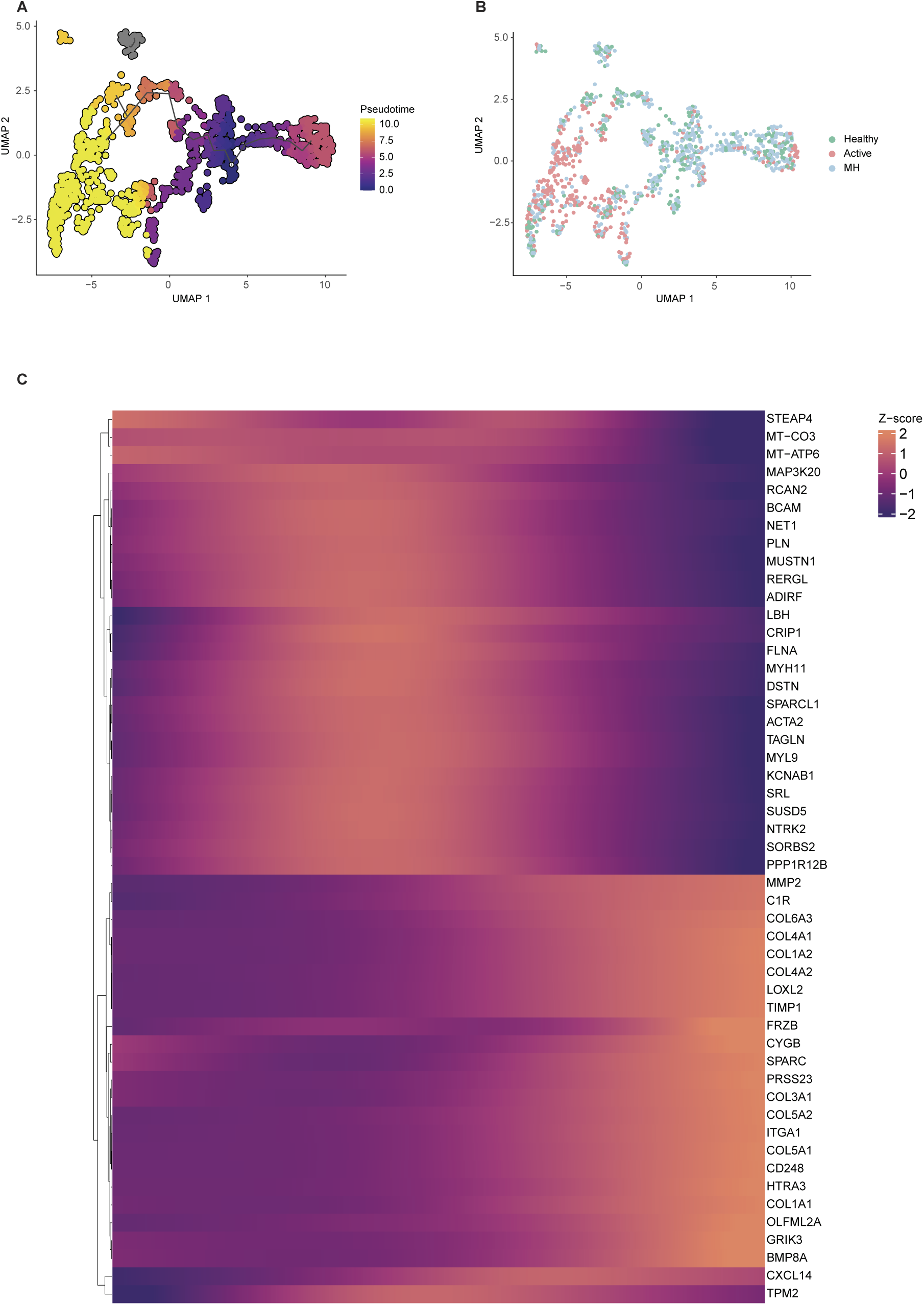
Pseudotime trajectory analysis indicates a transition of pericytes toward a matrix-remodeling, mesenchymal-like state. **(A)** Pseudotime trajectory of pericytes inferred using Monocle3 and projected onto the UMAP. The black line represents the principal graph connecting inferred cell states. Cells are colored by pseudotime. **(B)** UMAP embedding of the pericyte cluster from the scRNAseq dataset, colored by disease state. **(C)** Heatmap of the 50 most dynamic genes across pseudotime (Moran’s I, q < 0.05). Cells are ordered by pseudotime; genes are clustered into early and late transcriptional modules based on Z-scored expression.

**Supplementary Figure 9.**
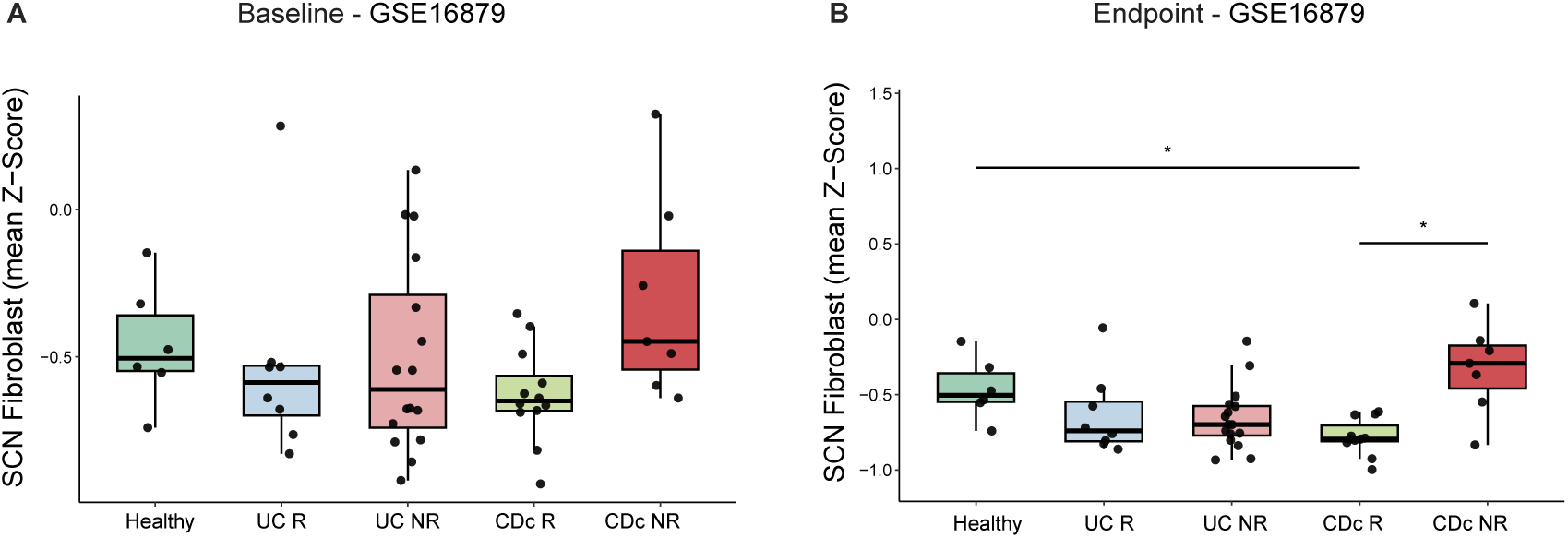
SCN fibroblast signature levels in IBD patients treated with Infliximab. (A) Boxplots displaying the baseline (pre-treatment) Z-scores of the SCN fibroblast signature in healthy controls, UC and colonic Crohn’s disease (CDc) patients from the GSE16879 cohort, stratified by subsequent clinical response to Infliximab (IFX). **(B)** Boxplots showing post-treatment Z-scores of the SCN fibroblast signature in the same cohort. Statistical significance was assessed using Wilcoxon rank-sum tests (*p < 0.05, **p < 0.01).

**Supplementary Figure 10.**
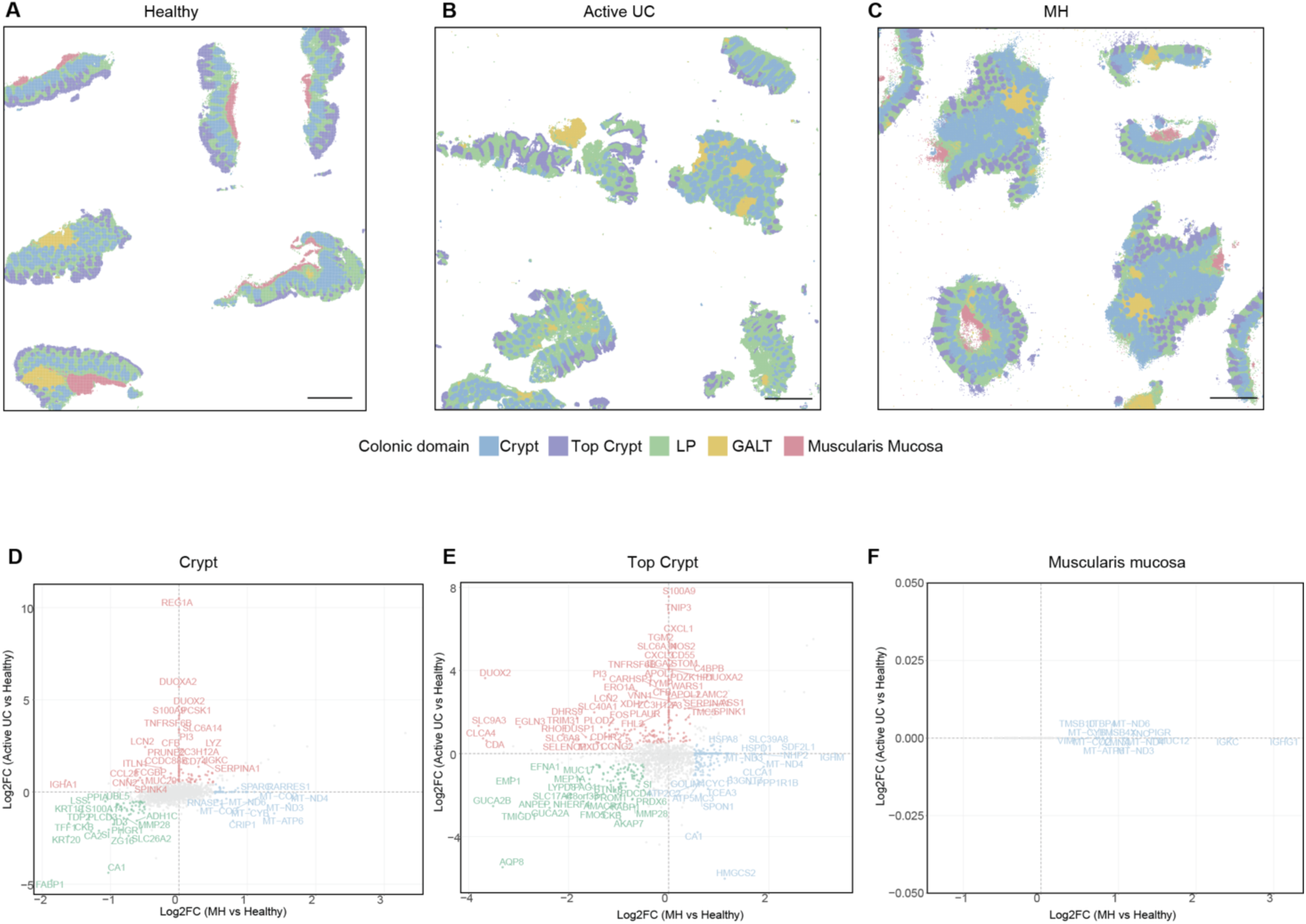
Visium HD spatial mapping and transcriptionally characterization of colonic domains. **(A-C)** Visium HD spatial domain maps analyzed using KNN clustering, colored by domain: Crypts (blue), Top Crypt (purple), Lamina Propria (LP, green), GALT (yellow), and Muscularis Mucosa (red). **(A)** healthy tissue **(B)** active UC **(C)** MH. Scale bar - 800µm. **(D-F)** Scatter plots showing domain-specific gene expression changes within the Crypt **(D)**, Top Crypt **(E)** and Muscularis Mucosa **(F).** Genes were categorized based on differential expression (|log2FC| > 0.5) and colored according to the disease state in which they are upregulated: healthy (green), active UC (red) and MH (blue).

